# Investigating microscopic angioarchitecture in the human visual cortex in 3D with angioMASH tissue clearing and labelling

**DOI:** 10.1101/2024.08.26.609648

**Authors:** Sven Hildebrand, Johannes Franz, Hanna Hoogen, Michael Capalbo, Philipp Bethge, Andreas Herrler, Fritjof Helmchen, Alard Roebroeck

**Affiliations:** Department of Cognitive Neuroscience, Faculty of Psychology & Neuroscience, Maastricht University, Maastricht, the Netherlands; Laboratory of Neural Circuit Dynamics, Brain Research Institute, University of Zürich, Zürich, Switzerland; Department of Anatomy & Embryology, Faculty of Health, Medicine & Life Science, Maastricht University, Maastricht, The Netherlands; Adaptive Brain Circuits in Development and Learning (AdaBD), University Research Priority Program (URPP), University of Zürich, Zürich, Switzerland; Neuroscience Center Zürich, Zürich, Switzerland

**Keywords:** Optical tissue clearing, Angioarchitecture, Light-sheet fluorescence microscopy, Neuroanatomy, Human visual cortex

## Abstract

Non-invasive imaging techniques, such as ultra-high field fMRI, are intricately connected to the underlying vasculature and are approaching ever higher resolutions. For the analysis of fMRI signals over cortical depth at such high resolutions, microvascular differences might have to be taken into account. Therefore, a better understanding of the laminar distribution and interareal differences in the cortical vasculature is becoming more important. However, in comparison to cyto- and myeloarchitecture, the study of angioarchitecture has received far less attention and relatively few methods have been described to visualise the vascular network in the human brain. Here we present angioMASH, a method for double labelling angioarchitecture and cytoarchitecture in archival human brain tissue, based on the recently published MASH protocol. The double labelling and optical clearing of thick human brain slices can be accomplished within 16 days. We use this method to acquire multi-resolution 3D datasets of combined cyto- and angioarchitecture in large (∼30 mm x 10 mm x 3 mm) volumes of human samples covering visual areas V1 and V2. We demonstrate for the first time, that classical angioarchitectonic features can be visualised in the human cortex and in 3D using tissue clearing and light-sheet microscopy. Lastly, we show differences in the vessel density and orientation over cortical depth within and between the two areas. Especially in V1, the vascular density is not homogeneous over cortical depth but shows distinct layering. These layers are also determined by changes in the orientation of the blood vessels from a predominantly radial to a more tangential distribution. In V2, differences in vascular density are less pronounced, but orientation profiles follow a similar trend over cortical depth. We discuss potential consequences of these differences for the interpretation of non-invasive functional imaging modalities such as fMRI or fNIRS.

**Graphical Abstract:** 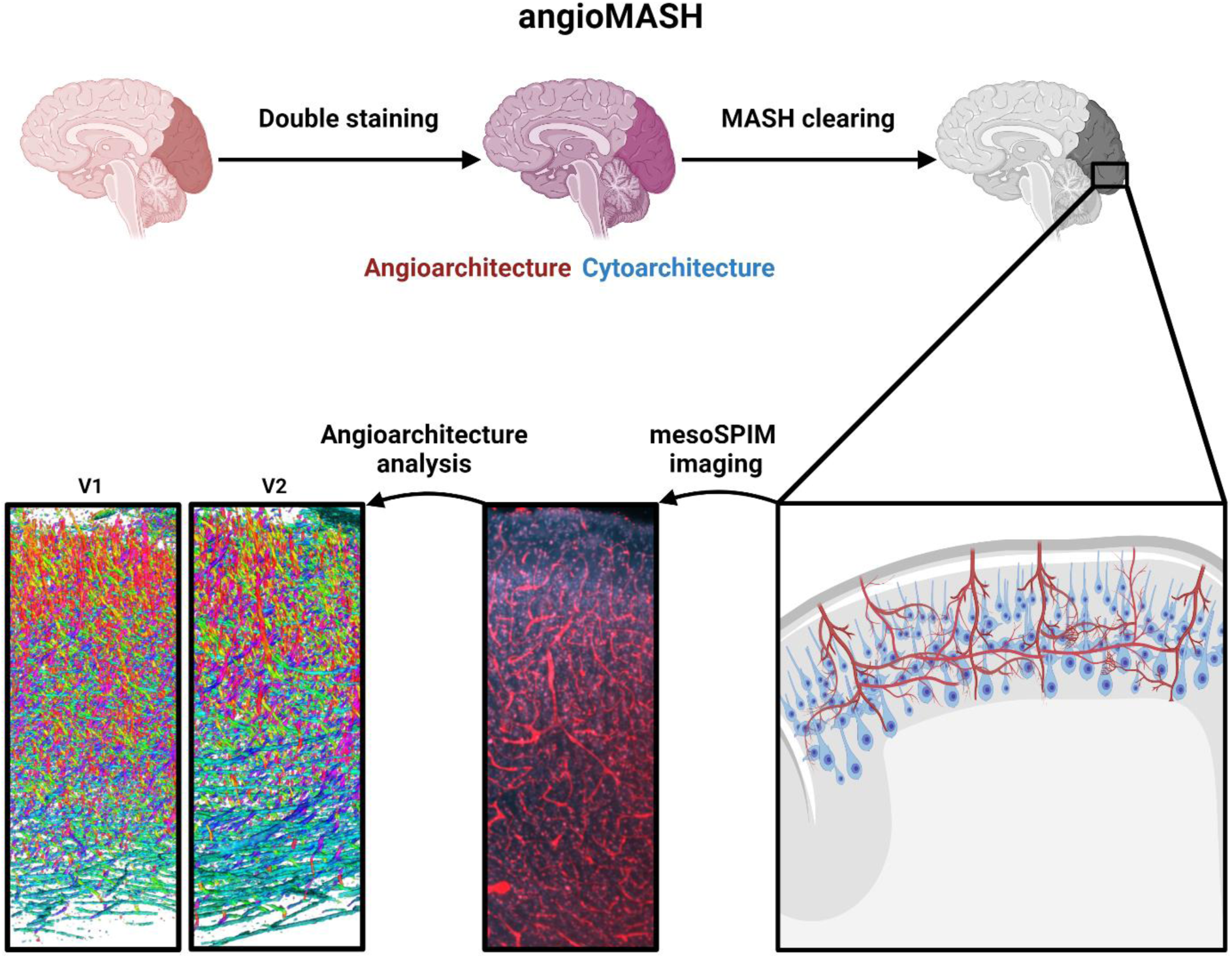

## 1. Introduction

The human brain has extremely high metabolic demands for its functioning^1^. In order to supply the nervous tissue with nutrients and oxygen, the brain contains an intricate vascular network of immense density and complexity. With the commencement of Pfeifer’s work over 90 years ago^2–4^ and later Duvernoy^5^, it has become clear that the human cortex is divided into horizontal vascular layers each characterised by a distinct density and distribution of arterioles, venules, and capillaries. This allows for the parcellation of the brain into areas with different angioarchitecture, homologous to the famous brain maps based on either cyto-^6,7^ or myeloarchitecture^8–14^. However, historically the angioarchitecture of the human cortex has received far less attention than the cyto- and myeloarchitecture, which are more closely related to the neuronal physiology and functioning of the cortex. Yet, the vasculature contributes indirectly but crucially to the brain’s function, supporting its metabolic needs, while the blood brain barrier protects the nervous tissue from harmful substances^15^. Understanding the variations of cortical vasculature in the human brain is becoming increasingly important. Not only are changes to the vasculature implicated in several neurodegenerative diseases, such as vascular dementia^16,17^ but the vascular network is also essential for non-invasive functional imaging modalities. Functional Near Infrared Spectroscopy (fNIRS)^18,19^ and functional Magnetic Resonance Imaging (fMRI)^20,21^ for instance, rely on the optical and magnetic properties of blood in cortical vasculature as the signal source. In recent years, technological advances, particularly at ultra-high magnetic field strengths, have enabled fMRI at resolutions high enough to distinguish both laminar and columnar mesoscopic structures^22–26^. Since the signal of both fNIRS and fMRI is only indirectly related to neuronal spiking and at the same time more directly linked to the brain’s blood flow, blood volume and oxidative metabolism^27,28^, it is important to understand the relations between cyto- and angioarchitecture. At the same time, the degree to which different cortical layers and areas as identified by cytoarchitecture correspond to variances in the organisation of angioarchitecture is not sufficiently understood. Furthermore, the vasculature itself seems to be organized in a laminar rather than columnar manner, which varies visibly between cortical areas^2–5,29^. Individual reports differ, but generally the highest capillary density can be observed in the lower part of the cytoarchitectural layer III and in layer IV, with the marked exception of primary visual cortex^5,30–32^. Such variations in vascular density across cortical depth are likely to result in similar variations of the fMRI signal sensitivity^33^. This problem might be addressable at the analysis stage but requires extensive knowledge about vascular density across cortical depth which is often missing and it is not clear to what extent these profiles are generalizable across individuals. Depending on the specific fMRI acquisition scheme, these depth profiles should reflect specific combinations of micro- and macro-vascular compartments. Further, the sensitivity of classical gradient-echo BOLD fMRI sequences is known to lead to poor specificity in the presence of large draining veins such that the signal recorded at superficial cortical depths also reflects functional activation at lower cortical depths. Given the heterogeneity of vasculature across brain areas, quantitative data on vessel densities and diameters across cortical depth per brain area might help to inform and extend the existing modelling and deconvolution approaches to eliminate these biases^34–37^. Finally, the fMRI signal has been shown to be sensitive to the relative orientation of the static magnetic field, to the orientation of the vessels, and this effect might also be dependent on cortical depth^38^. Quantitative data on the dominant vascular orientation, as well as its variation across brain areas and cortical depth might also enable modelling approaches in order to mitigate these biases at the analysis stage. Aside from the extensive, albeit only descriptive work of the early pioneers^2,3,39^, few studies have been conducted on human brain tissue so far^30,40^. A better understanding of the underlying anatomy could provide useful information to an imaging community aiming for ever higher field strength and resolution with fMRI techniques.

Recent developments in histological tissue processing and microscopy, enable fast volumetric imaging over large samples in order to address these questions in an unparalleled manner. Optical tissue clearing has become a widely used technique in many neuroscience labs, particularly for studies in rodents, as well as in other biomedical fields. Although initially its application to human brain tissue proved to be difficult because of its high lipid content, considerable progress has been made in recent years^41–46^, with solvent-based approaches showing especially high clearing capacity^41,44,46^. Therefore, rendering the tissue transparent is no longer the main challenge in the utilisation of optical tissue clearing in human brain tissue and the focus has shifted towards more sophisticated labelling strategies^47,48^ and data analysis^49–52^. Recently several publications have used lectins as labels for vasculature in combination with optical tissue clearing. So far however, this has been applied mainly in the mouse brain or on relatively small and thin human brain pieces^43,47,49,52,53^. Crucially, the ultimate goal would be to relate 3D microscopy imaging data to non-invasive, *in vivo* functional imaging methods. As these non-invasive methods have lower resolutions but can cover large field of views (FOVs), a scalable 3D histology method is needed that allows for FOVs approaching those of the functional methods.

Here we present angioMASH: a pipeline double staining angioarchitecture and cytoarchitecture in large optically cleared human samples. Our newly developed protocol combines the scalable, economic cytoarchitecture staining as well as the effective optical tissue clearing of MASH^41^ with protocols for labelling vasculature. The entire tissue processing takes 16 days to complete, from slicing the tissue to obtaining the transparent sample ready for microscopy. We use a combination of MASH, the SWITCH labelling approach^47^ and the protocol published by Harrison et al.^53^, to stain cortical angioarchitecture down to the capillary level in large human brain samples. We set out to investigate, whether angioMASH can visualise known vascular features such as the proposed layers of Duvernoy^5^ and then go beyond those descriptive results by quantitatively investigating vascular orientation distributions. We then compared the vasculature between visual area V1 and V2 and show marked differences in their density and orientation distribution over cortical depth. Finally, we stratified cortical depth-dependent distributions of vascular density by vascular thickness and provide summary statistics of these vascular features in the context of cytoarchitectonic layers and cell distributions.

## 2. Materials and Methods

### 2.1 Human brain tissue

Brain tissue samples were taken from three different healthy human body donors (see supplementary table 1 for donor data) of the body donation program of the Department of Anatomy and Embryology, Maastricht University. The tissue donors gave their informed and written consent to the donation of their body for teaching and research purposes as regulated by the Dutch law for the use of human remains for scientific research and education (“Wet op de Lijkbezorging”). Accordingly, a handwritten and signed codicil from the donor posed when still alive and well, is kept at the Department of Anatomy and Embryology Faculty of Health, Medicine and Life Sciences, Maastricht University, Maastricht, The Netherlands.

Brains were first fixed *in situ* by full body perfusion via the femoral artery. Under a pressure of 0.2 bar, the body was perfused by 10 l fixation fluid (1.8 % (v/v) formaldehyde, 20% ethanol, 8.4% glycerine in water) within 1.5-2 hours. Thereafter the body was preserved at least 4 weeks submersed in the same fluid for post-fixation. Subsequently, brains were recovered by calvarial dissection and stored in 4% paraformaldehyde in 0.1 M phosphate buffered saline (PBS) for 14-30 months.

### 2.2 Clearing and labelling of human brain samples

Each of the occipital lobes was cut into approx. 3 mm thick coronal slices. Of each of the three occipital lobes, three consecutive slices were further blocked into pieces of about 3×2 cm (see Suppl. Fig. 1-3).

Tissue blocks were then processed with angioMASH (Fig. 1), an adapted version of the MASH protocol^41^. We combined lectin labelling for vasculature as shown by Harrison et al.^53^ with the SWITCH labelling strategy^47^ and MASH^41^. The combination with SWITCH and performing sample delipidation before staining, rather than at the end, were used to improve label penetration in very large, thick human brain samples. The samples were kept on a shaker during incubation at all times. They were placed in 6 well cell culture plates and dehydrated 1h each in 5 ml 20, 40, 60, 80, 100% methanol (MeOH)/H_2_O (v/v) at room temperature (RT) and 1h in 100% MeOH at 4 ⁰C. Samples were then bleached overnight in freshly prepared, chilled 5% H_2_O_2_ in MeOH (v/v) at 4 ⁰C. This was followed by a delipidation step. For this, samples were placed in 50 ml tubes made of polypropylene as cell culture plates are incompatible with dichloromethane (DCM). Samples were incubated for 8h in 50 ml of 66% DCM/33% MeOH and 2x 1h in 100% DCM at RT. Samples were then stored overnight in 80% MeOH and rehydrated the next day in 5 ml each of 60, 40, 20% MeOH/H_2_O (v/v) at RT in 6 well cell culture plates for 1h respectively. This was followed by a second bleaching step in 50% (w/v) aqueous potassium disulfite solution for 1h at RT. The solution was stirred at approx. 70 ⁰C until all precipitated crystals were dissolved and filtered right before use. The solution can be reused several times. Samples were thoroughly rinsed 5x in distilled water and washed for another hour in distilled water at RT. Finally, samples were permeabilized 2x 1 h in 0.2% PBST (phosphate buffered saline + 0.2% (v/v) Triton x-100 adjusted to pH 7.4) and equilibrated overnight in SWITCH-OFF buffer (10 mM SDS in PBS at pH 7.4).

**Figure 1:**
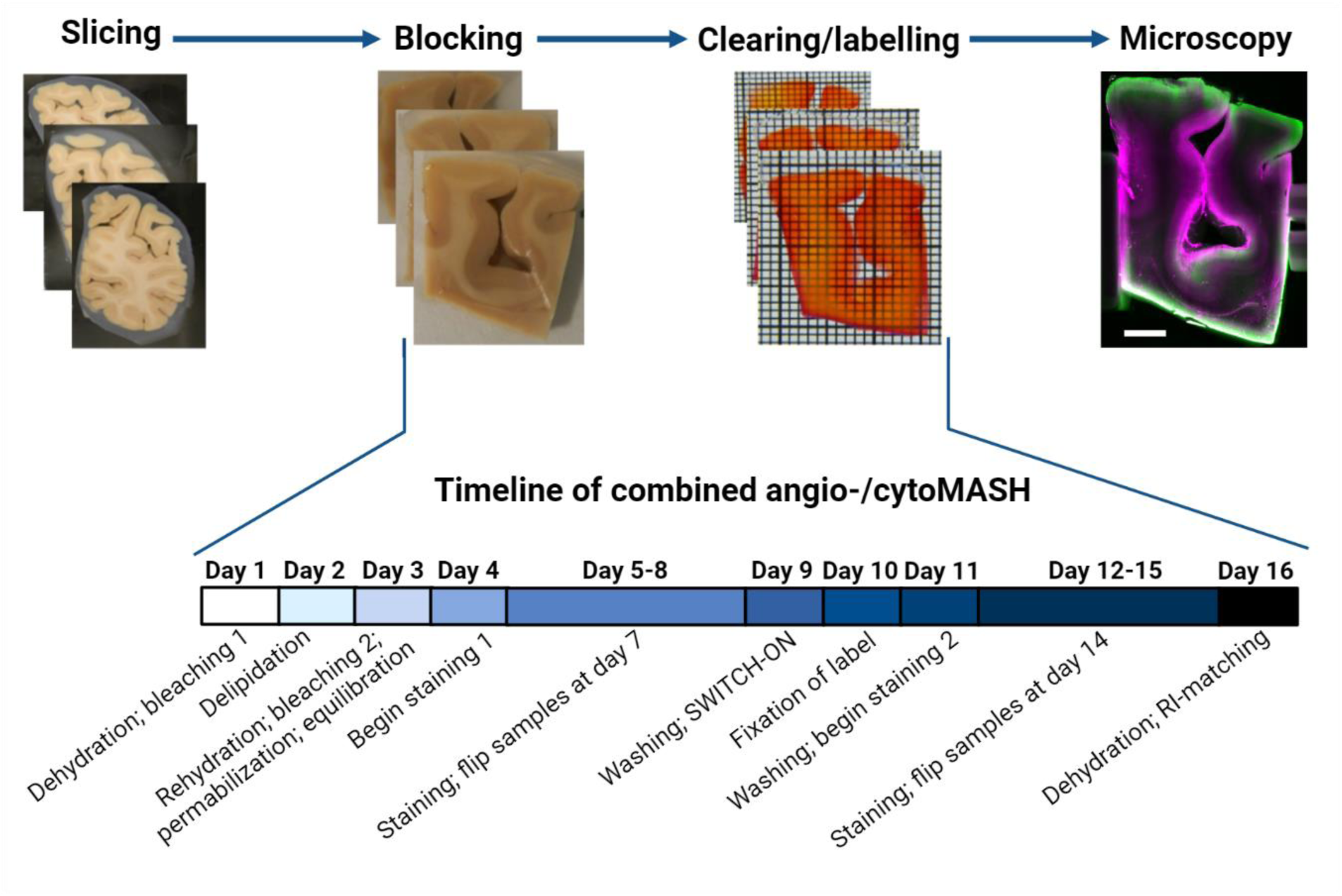
Workflow of the combined angio/cytoMASH pipeline. Top shows representative examples of the tissue at all major steps of the pipeline, from cutting coronal sections of the agar-embedded occipital lobes and blocking ROIs around the calcarine sulcus, to clearing and imaging. The detailed timeline for clearing and labelling of the tissue is given below. Scale bar: 3 mm; Grid 1×1 mm.

Samples were stained for angioarchitecture in SWITCH-OFF buffer containing *Lycopersicon esculentum* (LEL) lectin conjugated to DyLight®649 (Vector Laboratories, Inc.) diluted 1:100. Samples were incubated for 5 days and flipped after half the incubation time. Samples were then washed 3x 1 h in 5 ml 0.2% PBST, placed again in 50 ml tubes and incubated for 24 h in the same solution (referred to as the SWITCH-ON step). Subsequently, samples were incubated for 24 h in 4% PFA. This step empirically showed to improve the labelling quality. It is assumed that the lectins are fixed by the PFA. This was followed by 3x 1 h washed in McIlvain buffer at pH 4 (see McIlvain, 1921 for details)^54^. For MASH-NR counterstaining^41^, samples were placed again in 6 well cell culture plates and incubated in 5 ml 0.001% neutral red solution in McIlvain buffer at pH 4 for a duration of 5 days at RT. Samples were flipped after half of the incubation time. Finally, samples were washed again 2x 1 h in 5 ml of McIlvain at pH 4 and dehydrated 1h each in 5 ml 20, 40, 60, 80, 2x 100% MeOH/H_2_O (v/v). To ensure that all MeOH is removed before refractive index (RI) matching, samples were placed once more in 50 ml tubes and incubated 1h in 66% DCM/33% MeOH and 100% DCM respectively. Then samples were immersed in ethyl cinnamate (ECi). The angioMASH workflow is depicted schematically in Figure 1. Note that samples containing only grey matter can be immersed in ECi directly after the last incubation in 100% MeOH. However, in larger samples the white matter, especially the deep white matter, can remain slightly opaque.

### 2.3 Light-sheet microscopy

The samples were imaged on mesoSPIM light-sheet microscopy systems^55^ at the UZH in Zurich, Switzerland. The microscope consists of a dual-sided excitation path using a fiber-coupled multiline laser combiner (405, 488, 515, 561, 594, 647 nm, Omicron SOLE-6) and a detection path comprising an Olympus MVX-10 zoom macroscope with a 1× objective (Olympus MVPLAPO 1×), a filter wheel (Ludl 96A350), and a scientific CMOS (sCMOS) camera (Hamamatsu Orca Flash 4.0 V3). This set-up allows for nearly isotropic resolution by applying axial scanning in which the beam waist is scanned with electrically tunable lenses (ETL, Optotune EL-16-40-5D-TC-L) synchronized with the rolling shutter of the sCMOS camera. The MASH-NR cytoarchitecture label (neutral red) was excited with 561 nm and DyLight®649-conjugated lectin with 647 nm. For both channels, a multiband emission filter (QuadLine Rejectionband ZET405/488/561/640, AHF) was used. We first acquired large FOV overview scans of all brain samples with dual sided illumination at a pixel size of 10.52 µm and a step size of 10 µm. The cytoarchitecture staining had lower wavelength excitation and emission spectra and hence could not be imaged as deeply as the far-red vessel label. We therefore acquired slightly higher resolution (6.55 µm in plane and 6 µm z-step size) overviews with single sided excitation by tilting the sample 20⁰ with respect to the excitation light. In this orientation, the excitation light has to travel through less tissue when imaging the more medial parts of the sample and, therefore, the illumination of the cytoarchitecture channel (shorter wavelength) was improved. We then acquired tiled high-resolution (2.03 µm pixel size and 2 µm z-step) data around the V1/V2 border, identified beforehand based on a macro-anatomical landmark, the stripe of Gennari. Finally, for one sample, we cut off a small gyrus containing parts of V1 and V2 on each side of it and acquired separate data sets at the highest resolution the mesoSPIM allowed (1.03 µm pixel size with 1 µm step size). We opted to cut off this part as the cortical wall was relatively straight and it allowed for an illumination perpendicular to the cortical layers, so that an illumination gradient over the cortical depth could be avoided.

### 2.4 Data analysis and visualisation

Image pre-processing was done in FIJI^56^. All overview images acquired with a 20⁰ tilt were resliced and rotated for better visualisation. Tiled high-resolution acquisitions were stitched using the BigStitcher plugin^57^. Misalignment of the cell- and vessel-images due to chromatic aberration was corrected using a rigid transform, applied with the interest-point-based registration available in BigSticher. Interest points were detected in both images, using a Difference of Gaussian filter, which was manually tuned to the average size of cells visible in the vessel image. For the orientation analysis, vessel segmentation and skeletonisation was carried out in FIJI^56^. To this end, the high-resolution raw datasets were first filtered with a 3D Laplacian of Gaussian filter using a FIJI plugin^58^. Data was separately filtered with a sigma of 2 and 6 pixels in each direction (2.06*2.06*2 µm and 6.55*6.55*6 µm, respectively), to better preserve capillaries in the case of the smaller filter setting and arterioles and venules in the case of the larger filter setting. Binary masks of these two filtered images were created by thresholding in FIJI and added together to generate one segmentation of the micro-vasculature. This binary dataset was used as the basis for the orientation and depth analysis, as well as for the skeletonisation used to extract vessel centrelines. Cytoarchitcture layers were delineated by two expert annotators. A consensus annotation was reached and in case of uncertainties (specifically with the boundaries of layer IVa in V1), the layer borders were indicated as tentative with dashed lines. Overall, the layer annotation was highly consistent over depth of the data set (see Suppl. Fig. 4).

#### 2.4.1 Vascular density

Large vessels were segmented separately. These vessels were oriented radially with respect to the cortex, and sometimes collapsed. In both cell and vessel images, these collapsed parts were characterized by lower auto-fluorescence than the surrounding tissue which allowed segmentation in a down-sampled (3.09*3.09*3 µm) dual-channel image using the LabKit plugin^59^, and manual clean-up using Napari^60^. The micro-vascular segmentation occasionally contained a few round, cell-like, objects, potentially visible as a consequence of auto-fluorescence. As these might influence the density analysis, these objects were filtered using an area-threshold for connected components implemented using dask-image and Python. The filtered micro- and the macro-vascular segmentations were then joined for the purpose of the density analysis.

Cells were segmented using StarDist^61^. Training data was generated semi-manually. A total of eight volumes of interest (VOIs), each 150*150*150 voxels, were selected as training data. The specific volumes were selected to reflect the variety seen in the full data in terms of signal-to-noise, cortical layer and cell morphology. Following background-removal and equalization as described below, cells were segmented using Labkit and segmentations were post-processed using ITK-SNAP^62^ and Napari. We followed the recommendations for training and application of a 3D StarDist model, using 96 rays, an anisotropy of (1, 2, 2) to account for the median object size of (13.5, 6, 6.5) observed in the training data, and distributed prediction for big images. Finally, cells whose central voxel was also labelled in the combined vascular segmentation were removed from the cell segmentation.

The cortical ribbon was segmented manually in ITK-Snap. To this end, the data was down-sampled to 2.06*2.06*10 µm, largely preserving in-plane resolution while taking only every 10^th^ image across depth. The data was then equalized in the case of the angio-channel or background-removed and equalized in the case of the cyto-channel. Fast equalization and background removal were done with adapted versions of ClearMap^63^. Because these pre-processing steps may introduce small artefacts, we only used them as a visualization aid in the process of segmentation. For obtaining a definition of cortical depth, we used in Laynii^64^. We labelled 1) the imaging medium near the cortical/pial surface, 2) the white matter (WM), and 3) missing data (e.g. damaged tissue) that could be excluded from downstream analysis. Gray matter (GM) was then inferred as the un-labeled tissue. Finally, we labeled regions of the image where GM-WM interface or the GM-pial-surface interface are just outside of the imaged volume. In regions that are closer to these missing reference points than to present reference points, cortical depth cannot be established unambiguously and these regions were excluded. The labelled image was then post-processed. First, the data labelled as excluded was dilated with a disc-shaped structuring element of 5 voxels, to conservatively preserve this data during the following steps. Then, each label was smoothed with a Gaussian kernel with a standard deviation of 5 voxels. Next, the label image was down-sampled again but only in plane to achieve an isotropic resolution of 10.3 µm. Finally, the labelled image was processed with Laynii in order to transform the GM segmentation to cortical depth coordinates. The tissue samples used here have been selected because of their location in relatively flat parts of the cortex. Therefore, in this data, cortical layers can be assumed equidistant throughout the tissue. Similarly, for the present dataset, cortical thickness stays similar throughout the analyzed volume. Therefore, it was possible to report depth profiles not only in terms of normalized cortical depth, but also in terms of absolute cortical depth.

#### 2.4.2 Vascular density over depth

Vascular density was calculated over cortical depth in 100 equi-distant histogram-bins, as the fraction of the number of voxels labelled in the combined micro- and macro-vascular segmentation relative to the total number of voxels, of each bin. This was implemented as a chunked computation, involving lazy up-sampling of the cortical depth metric to the native image resolution, and efficient histogramming. Specifically, for the present analysis, the image data and coordinate system were chunked such that 10 consecutive in-plane images formed one chunk. This made it convenient to obtain standard errors of the mean for the density estimates and to assess the variability of results that could be expected from traditional 2D imaging of histological sections.

Cell densities were extracted using the same analysis as presented above. Unlike vessel density, which we present as a fraction of the imaged volume, cell densities are reported as an average cell-count per cubic millimetre.

In addition to overall vascular density, we also analysed vascular density stratified by the local thickness of vessels. For this, we used the fast, approximate algorithm presented by Dahl^65^.

#### 2.4.3 Orientation analysis

For the orientation analysis of the vasculature, we segmented the images into 1) the four angio-architectonic layers proposed by Duvernoy^5^, and 2) a three-compartment model based on the cyto-architecture. Coordinates of lines defining the layer borders were selected manually in Fiji using maximum intensity projections (MIPs) over successive bins of the binarized data. We choose not to delineate all six cytoarchitectonic layers, since the aim was to use the functional compartmentalization often employed by the layer-fMRI community. Therefore, the broader classification of supragranular, granular, and infragranular layers was chosen. This was done because it is often assumed that these compartments sub-serve different functions and underlie different connectivity within models of the canonical cortical microcircuit^66–68^. The larger size of these compartments also makes them more comparable to UHF-fMRI data. These three layers are referred to here as “functional compartments”, to distinguish them from the standard six cytoarchitectonic layers. The selection of the functional compartments was done with the raw data of the cell body staining as an overlay. The two sets of layer coordinates were selected for each staining, one based on MIP over the first 100 µm (angioarchitecure) and 30 µm (functional compartments) of the 1mm thick stack and one over the bottom 100 µm (angioarchitecture) and 30 µm (functional compartments). MIPs were created over these different bin-widths to allow for the best visual delineation of the layers for each channel. To delineate the layer boundaries as accurately as possible, we generated a ‘surface’ through the stack by linearly interpolating between the drawn lines at the top and the bottom of the stack. This yielded one line delineating the layers for each individual image. Based on the upper and lower boundary of each layer, we then calculated a line in the middle of each layer. This middle line was later used for layers straightening. This way all lines in all images follow the curvature of the layer. Layer coordinates were then used to segment the binarized image stack into four (angioarchitectural layers) or three (functional compartments) stacks, which only contained one layer each. These layer stacks were then straightened using the Fiji “Straighten” function, using the lines defined earlier. Orientation analysis was conducted both on the unstraightened and the straightened layer stacks. The orientation analysis was performed with the Fiji Plugin OrientationJ^69,70^ with a local window of 8 pixels. The OrientationJ output consists of an absolute frequency distribution of orientations of all pixels, for each image of the stack. Means and standard deviations over all images and for each orientation were extracted and plotted using Python. OrientationJ also generated the colour-coded orientation maps with the HSB Color Survey function for each image in the stack. These colour maps were visualised in 3D with Vision4D (v4.0, arivis AG, Munich, Germany).

#### 2.4.4 Orientation over depth

Finally, we also analysed vessel orientation across cortical depth. Depth-profiles were extracted using the same analysis as presented above (see 2.4.2). Note that, in contrast to the analysis of straightened vascular compartments presented above, this analysis was run on the raw orientation maps. That is, cortical curvature might affect the results of this analysis. However, given that the present data was selected because the cortex shows relatively little curvature, we believe this effect to be small.

#### 2.4.5 Visualisation

Figures have been created using FIJI, matplotlib, ParaView, and Biorender.com. Large parts of the analysis were scripted in Python as well as the Fiji macro framework and are openly available at https://github.com/ivychad/Vasculature-Analysis-Pipeline-Code and https://github.com/jojofranz/pydepthmap.

## 3. Results

We report the results in five parts. In section 3.1, we report the extension of the MASH clearing and labelling method to the combined investigation of cytoarchitecture and angioarchitecture in the human brain. In section 3.2, we demonstrate neuroanatomical intra- and inter areal differences of angioarchitecture in V1 and V2 with this new pipeline. Subsequently, we quantify vascular and cell density profiles across cortical depth in section 3.3, and vessel orientation profiles across angioarchitectural layers in section 3.4. Finally, we quantitatively compare vessel orientation profiles across infra-, supra- and granular compartments.

### 3.1 Combined investigation of cortical vasculature and cytoarchitecture in large, thick human brain samples

We could successfully combine our existing MASH clearing and cytoarchitecture labelling protocol with deep vascular staining. The angioMASH approach (Fig. 1) allows for a deep penetration of the lectin label over several millimetres of tissue (Fig. 2 and Suppl. Video 1). Even at relatively low resolutions, marked changes in the angioarchitecture between V1 and V2 are visible, in particular the vessel dense stripe at (cytoarchitectural) layer IVb-c in V1 (see arrows Fig. 2). This matches the description by earlier authors^5,29^, who describe the internal granular layer of the striate cortex as possessing the densest vascular network in the entre brain.

**Figure 2:**
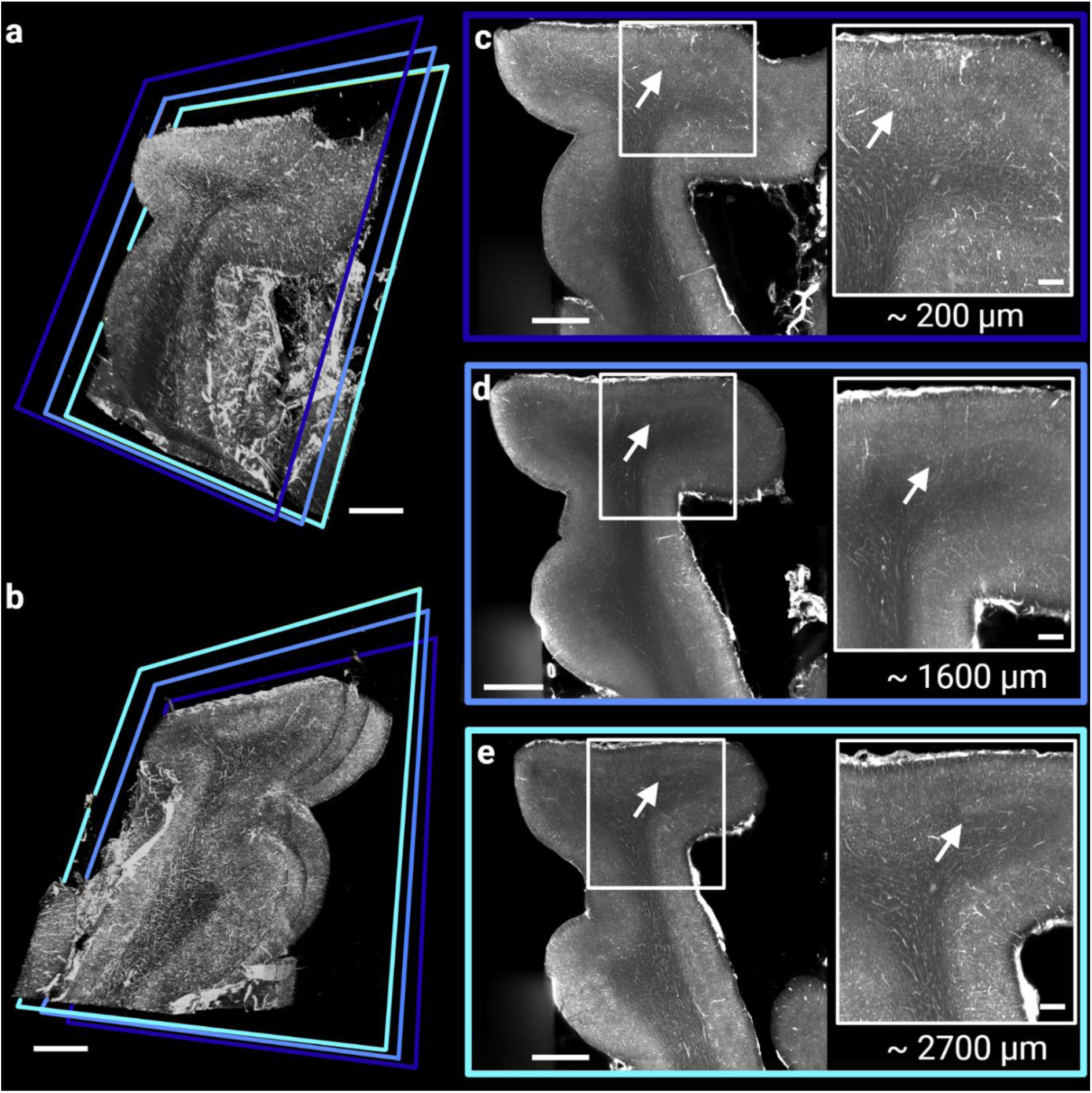
Overview acquisitions of the samples illustrate the quality and depth penetration of the vasculature staining. 3D rendering of a sample taken from Occipital lobe 3, sample 2, showing the posterior (**a**) and anterior (**b**) side. The frames indicate the approximate positions of the MIPs shown in **c**-**e**. Panels **c**-**e** were projected over 150 µm located at the posterior side (**c**; ∼ 200 µm deep into the tissue), middle (**d**; ∼ 1600 µm deep into the tissue), and anterior side (**e**; ∼ 2700 µm deep into the tissue) of the sample respectively. The vascular label is clearly visible throughout the whole sample thickness with a particularly dense (bright) stripe demarcating V1 (stopping at, and slightly above, the white arrows). All panels show the data in inverted greyscale. Scale bars: 2 mm for each full view and 500 µm for each magnified panel.

Multi-scale data sets could be acquired with an isotropic resolution of ∼10 µm to 2 µm for all the samples. A representative example of the different resolution levels is shown in Figure 3. The light scattering in large FOV overview scans is very strong towards the medial sample regions (towards the deeper cortical layers and white matter, when illuminating the tissue from the pial surface), especially in the lower wavelength cytoarchitecture staining (Fig. 3a: excitation light coming from the top and bottom of the panel). Therefore, higher resolution scans where performed, which still provided an overview over a large part of the sample and in which the cytoarchitecture staining could be better appreciated (Fig. 3b). In these acquisitions, the sample was rotated 20° oblique to the direction of the exciting light. For each sample, multiple tiles were acquired at high-resolution covering an extensive area around the V1/V2 border (Fig. 3c and Suppl. Video 2). Figure 3d shows a zoom-in of the highest resolution of 2×2×2 µm acquired for all samples, illustrating qualitatively the capacity to distinguish microscopic elements of both cyto- and angio-architecture.

**Figure 3:**
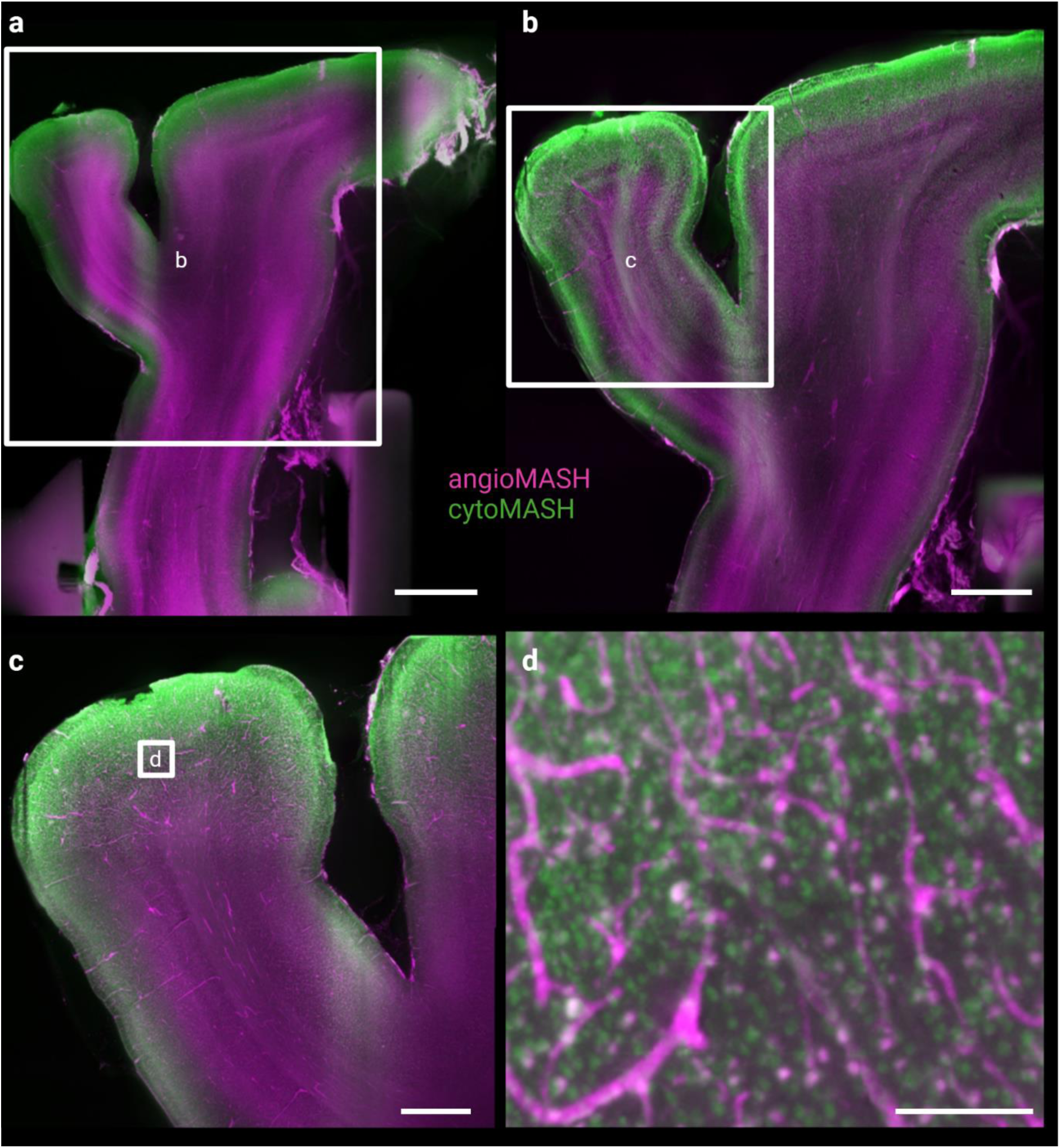
Representative example of the multi-scale data. **a)** We first acquired fast overview scans with the highest FOV possible and a voxel size of about 10 µm (MIP over 50 µm; scale bar: 4 mm) showing the cytoarchitecture stain (cytoMASH) in green and the vessel label (angioMASH) in magenta. **b)** Light scattering is especially prominent in very large tissue pieces combined with a classical light-sheet geometry, in which the excitation light has to travel through the entire lateral length of the sample. We therefore performed overview acquisitions at slightly higher resolution (6 µm) as well, in which the sample was rotated 20° (MIP over 60 µm; scale bar: 2 mm). **c)** Tiled and stitched high-resolution dataset of the ROI around the V1/V2 border (MIP over 50 µm; scale bar: 1 mm). **d)** Magnified region indicated in panel c, demonstrating the resolution sufficient to distinguish individual capillaries and cells (scale bar: 100 µm).

It can be appreciated that the choice of colour channels was chosen to maximize imaging quality for the angioarchitecture, i.e. longer, more deeply penetrating wavelength for the angio-channel and shorter, less deeply penetrating (and therefore more scattering) wavelength for the cyto-channel. In these datasets, individual cells and capillaries can be clearly distinguished (Fig. 3d).

### 3.2 Neuroanatomical intra- and inter areal differences of angioarchitecture in V1 and V2

In order to allow for the best comparison between V1 and V2, two separate datasets were acquired at the highest resolution that the system allowed for (1×1×1 µm). The illumination direction for these two datasets was perpendicular to the cortical wall and hence parallel to the layers, which allowed for better inspection of the architecture. To enable this illumination direction, the sample had to be physically cut, perpendicular to the cortical wall, in a region where V1 and V2 were located opposite to each other. As seen in the higher FOV datasets, the shorter wavelength of the excitation light in the cyto-channel (561 nm vs 647 nm for the angio-channel) leads to more light-scattering and hence a gradient in the image from left to right. A qualitative comparison of the angioarchitecture between V1 and V2 in relation to the cytoarchitectural layers reveals striking differences, both in the overall density as well as the distribution of the vasculature (Fig. 4). For best visualisation of both features, the MIPs for the angio- and cyto-channels were created over different depths. Angioarchitecture was judged to be best visible at projections over at least 100 µm, whereas cytoarchitecture was not ideally visible in such thick projections. We opted therefore for 150 µm MIPs for angioarchitecture and 30 µm MIPs, taken from the middle, for cytoarchitecture. In V1, the vascular layers appear more pronounced in general. Blood vessels are mainly radially organized in supragranular layers and many larger vessels terminate in layer III. The orientation of the blood vessels is more diverse in layers IVa-b, and a particularly dense mid-cortical vascular layer is visible in the lower part of layer IVb and within layer IVc. The main vessel orientation changes abruptly in the infragranular layers, where they run mainly tangentially. The border between grey and white matter is clearly visibly by a sharp decline in vascular density, while retaining mainly tangential orientation. In both V1 and V2, two distinct vascular sub-layers are visible within cytoarchitectural layer I: The first sub-layer is almost devoid of small capillaries while the second one shows a marked increase in vessel density, caused mainly by smaller vessels. In V2, it was particularly difficult to determine a visible difference between Duvernoy’s layers Ib and II. In general, the layers were less striking in this area. The border between layers II and III is marked by the termination of many smaller radial vessels, although this border is not as visible as compared to V1 and the delineation is more tentative. The seemingly densest layer, layer III, extends roughly to the borders of the cytoarchitecturally defined layer IV. After that, vessel density decreases and the general orientation of the vasculature seems again to be predominantly tangential.

**Figure 4:**
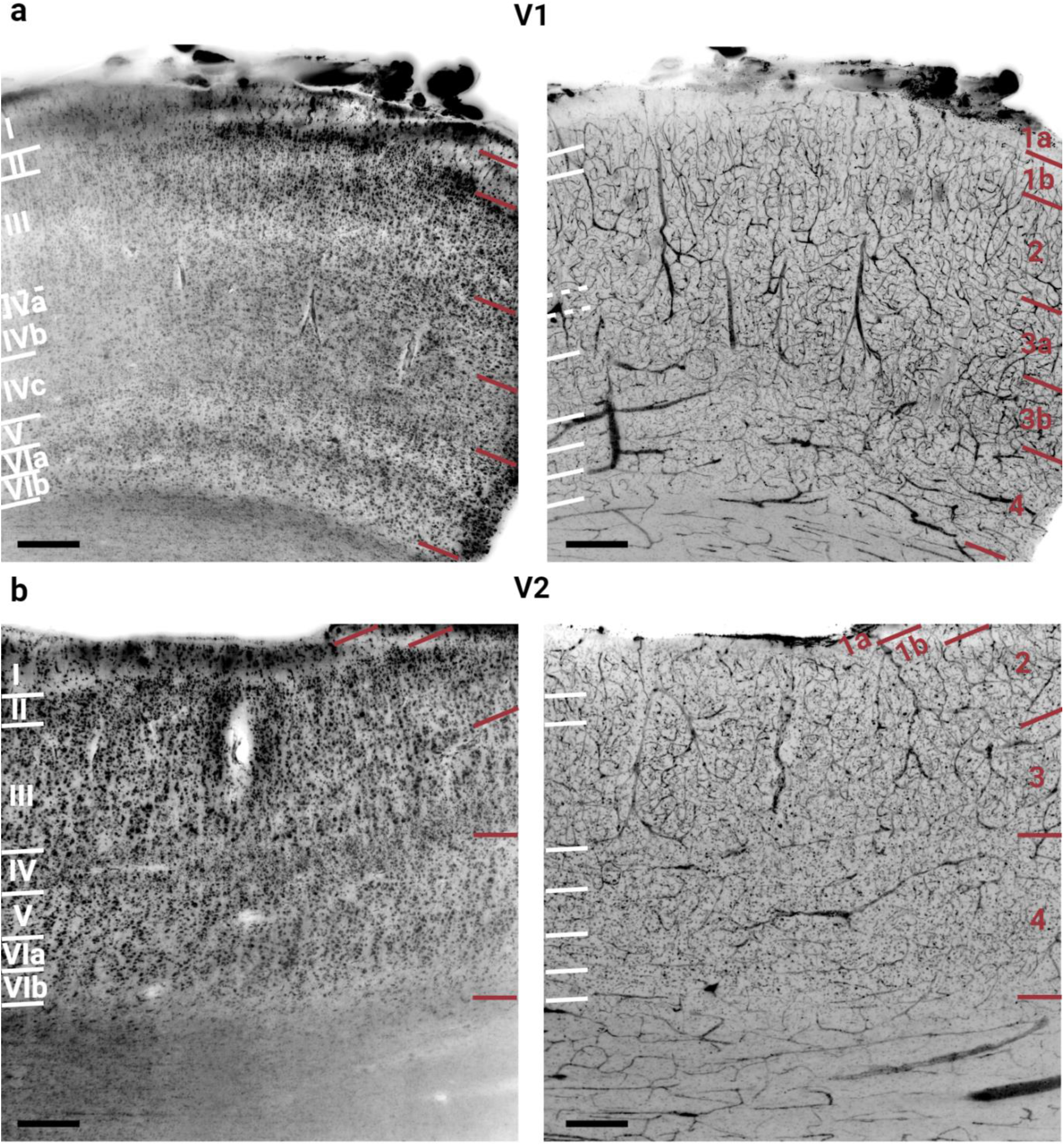
Joint visualisation of angio- and cytoarchitectural layers of V1 and V2. **a)** Cytoarchitecture (left) and angioarchitecture (right) of primary visual cortex (V1). **b)** Cytoarchitecture (left) and angioarchitecture (right) of secondary visual cortex (V2). Layers for angioarchitecture were based on the description of the 4 principal layers according to Duvernoy5 (red labels). The layers as identified by cytoarchitecture (white labels, based on the description by von Economo and Koskinas7) are indicated in the angioarchitecture panels as well and vice versa. Note that the boundaries of layer IVa in V1 were not clearly definable and are indicated tentatively with dashed lines. The MIPs were generated over 150 µm for angioarchitecture and 30 µm, taken from the middle of these 150 µm, for cytoarchitecture. Scale bars: 250 µm respectively.

### 3.3 Vascular and cell density profiles across cortical depth

The comparison of cell densities over cortical depth reveals a higher overall cell density for V1 compared to V2 (Fig. 5a). These differences are particularly pronounced in lower superficial layers as well as in layer IVc and infragranular layers (compare Fig. 5a, and green lines with layer indications in c). Over relative cortical depth, the internal granular layer IV of V2 coincides with the relatively cell poor layer IVb of V1, resulting in similar cell densities in that region. Overall, the higher cell densities in V1 vs V2 as well as the profiles across depth match well with previous observations by other authors^7^. Additionally, V1 demonstrates an overall much higher vessel density than V2 as well (Fig. 5b). This observation is well in line with findings from other researchers in both human^5^ and non-human primate^29,32^. Both areas show a low increase in vascular density towards the middle to lower third of the cortical thickness, followed by a steady decrease of vascular density in the infragranular layers and towards the white matter (see Fig. 5b and magenta lines with layer annotations in c).

**Figure 5:**
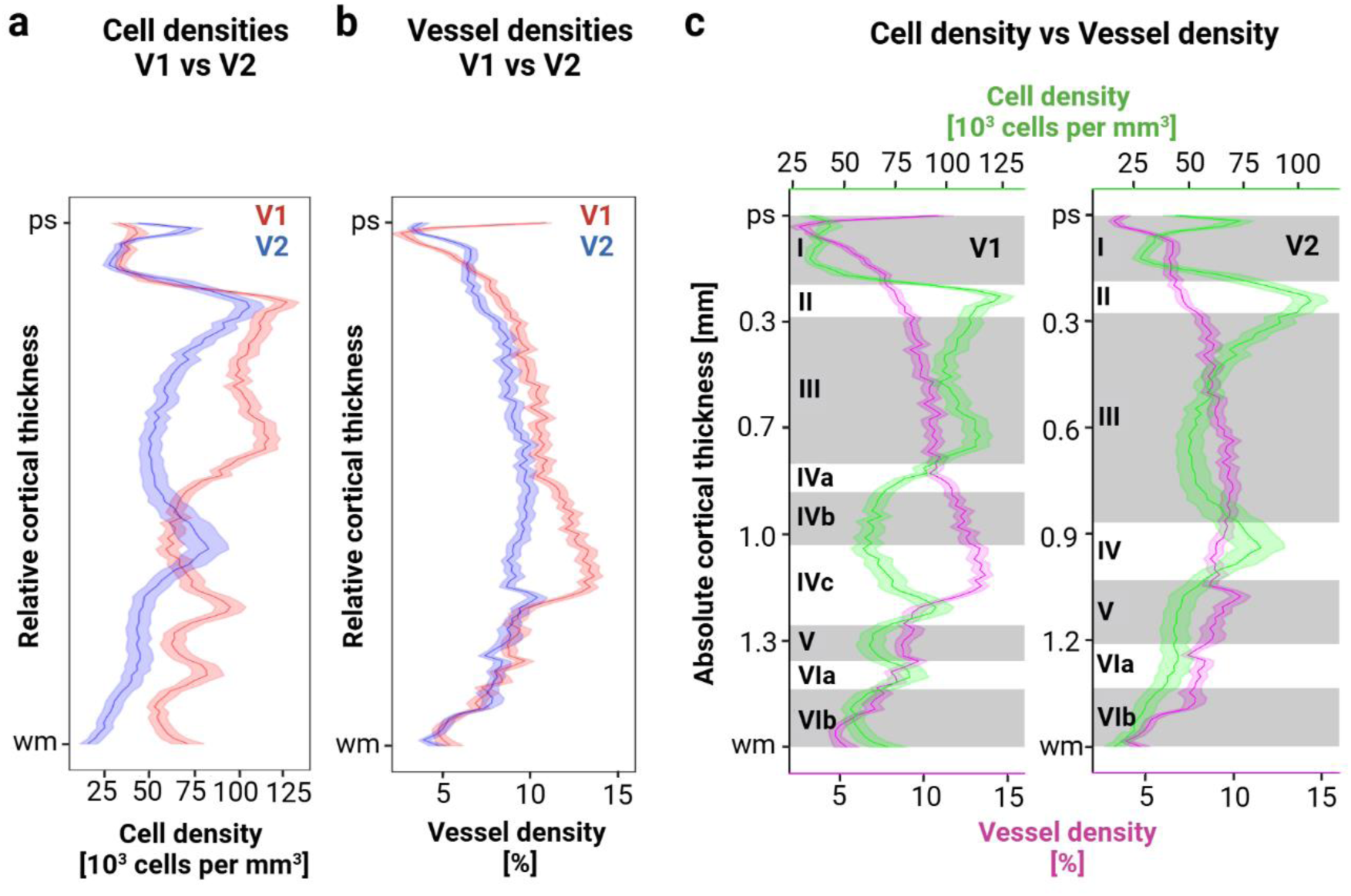
Density profiles of both vasculature and cells for V1 and V2. **a**) Comparison of cell densities as number of cells per mm^3^ along relative cortical depth in V1 (red) and V2 (blue). **b**) Comparison of vascular densities, displayed as the percentage of segmented pixels per histogram bin, along relative cortical depth in V1 (red) and V2 (blue). **c**) Comparison of cell densities (green) and vessel densities (magenta) across absolute cortical thickness within V1 (left) and V2 (right). Approximate cytoarchitectonic layer indication are provided in roman numerals in the left side of each diagram. Solid lines show the mean density, lighter shaded areas ±2x the SEM.

The highest vascular density in V1 occurs within cytoarchitectural layers IVb and IVc. In V2 in contrast, there is no clear peak in vessel density visible in the granular layer IV. The small peak in layer V is likely explained by one particularly large vessel branching within and crossing through this location in the 3D volume (Fig. 6). The few large blood vessels cannot however account for the overall vessel density, which seems to be mainly dictated by the amount of microvasculature. Overall, the density comparison shows that there is higher vascular density in the supragranular and granular layers in V1 as compared to V2, while the infragranular vessel density is similar for both areas.

**Figure 6:**
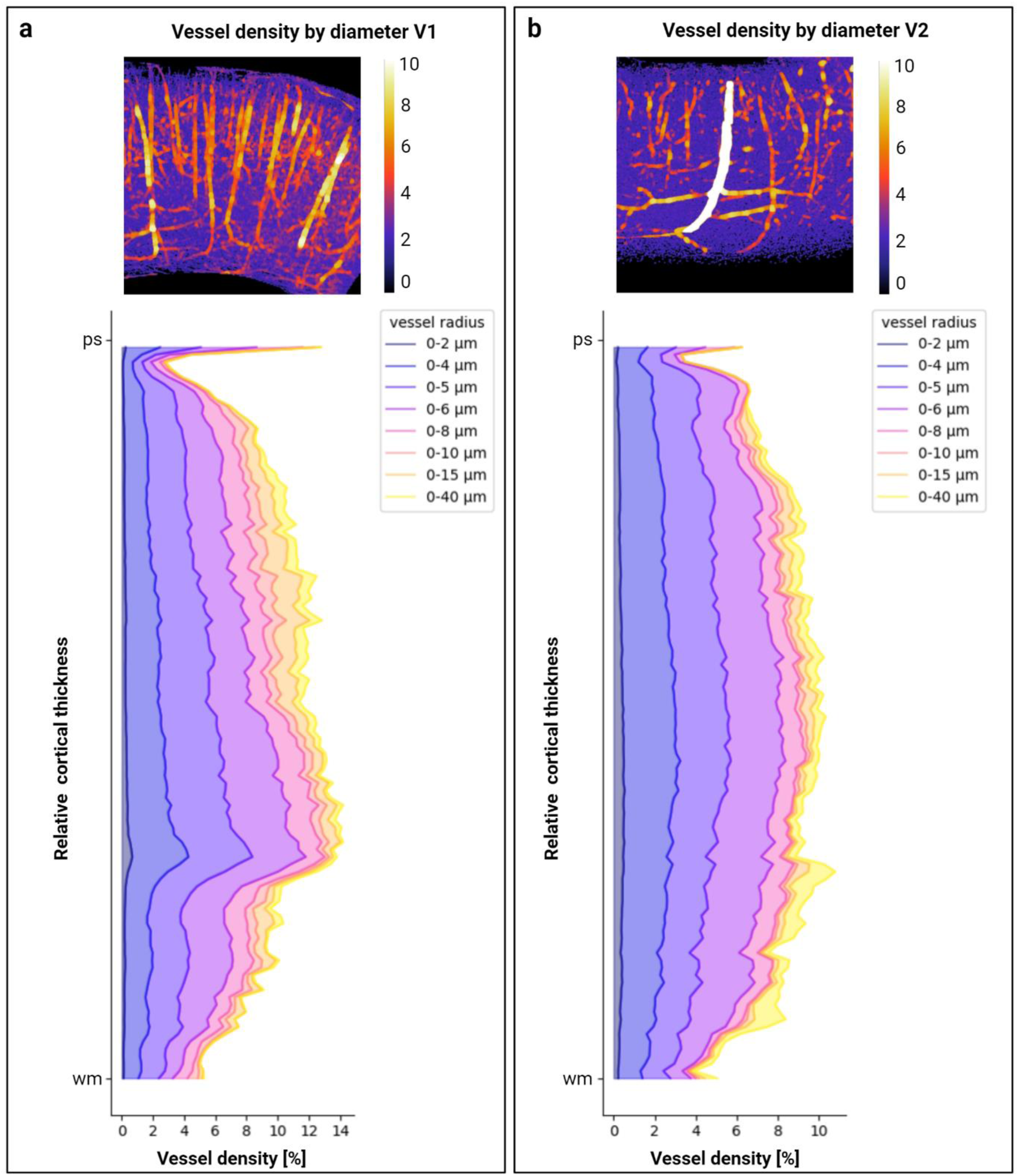
Vascular density across cortical depth stratified by vessel thickness. **a**) Percentage of vessel density per image plane of the V1 dataset stratified into 8 bins by the diameter of the segmented blood vessels (blue: smallest vessels, yellow: largest vessels). **b**) Percentage of vessel density per image plane of the V2 dataset stratified into 8 bins by the diameter of the segmented blood vessels (blue: smallest vessels, yellow: largest vessels).

### 3.4 Vessel orientation profiles across angioarchitectural layers

In order to investigate the qualitatively observed vascular features in a more objective manner, the vessel orientations were analysed between the different angioarchitectural layers in V1 and V2 (Fig. 7). For this, the four main angioarchitecture layers, as identified by the layering scheme of Duvernoy^5^, were manually segmented in the entire stack, for both V1 and V2 (see Suppl. Fig. 5 and 6). Despite imaging on the sulcal wall, rather than the fundus or gyral crown, both datasets have a noticeable natural curvature of the cortex ribbon, both in-plane as well as through-plane. To account for this curvature, we show the results of the orientation analysis of angioarchitectural layers after straightening the manually segmented compartments (see section *2.4.1* *Orientation analysis*). V1 shows more blood vessels with a radial orientation in the two more superficial layers 1 and 2 (Fig. 6: -90° and 90° (red) correspond to a vertical or radial orientation, 0° (cyan) to a horizontal or tangential orientation), which is in agreement with the qualitative description above (section 3.2; Fig. 4) and by other authors^4,5,32^. Layer 1 (and layer 2 to a much lesser degree) also show a smaller second peak around the tangential orientation. Both lower layers 3 and 4 of V1 show more tangentially oriented blood vessels and both curves are almost congruent in their orientation profiles. In area V2 the orientation profiles are more evenly distributed across the different layers. There is still a tendency for more tangential blood vessels in the deepest vascular layer 4 and, to lesser extent, more radially oriented vessels in layers 1 and 2. These observations do not change fundamentally in the curved, unstraightened data, except that the orientations show a wider distribution in every layer and for both areas, as would be expected (Suppl. Fig. 6).

**Figure 7:**
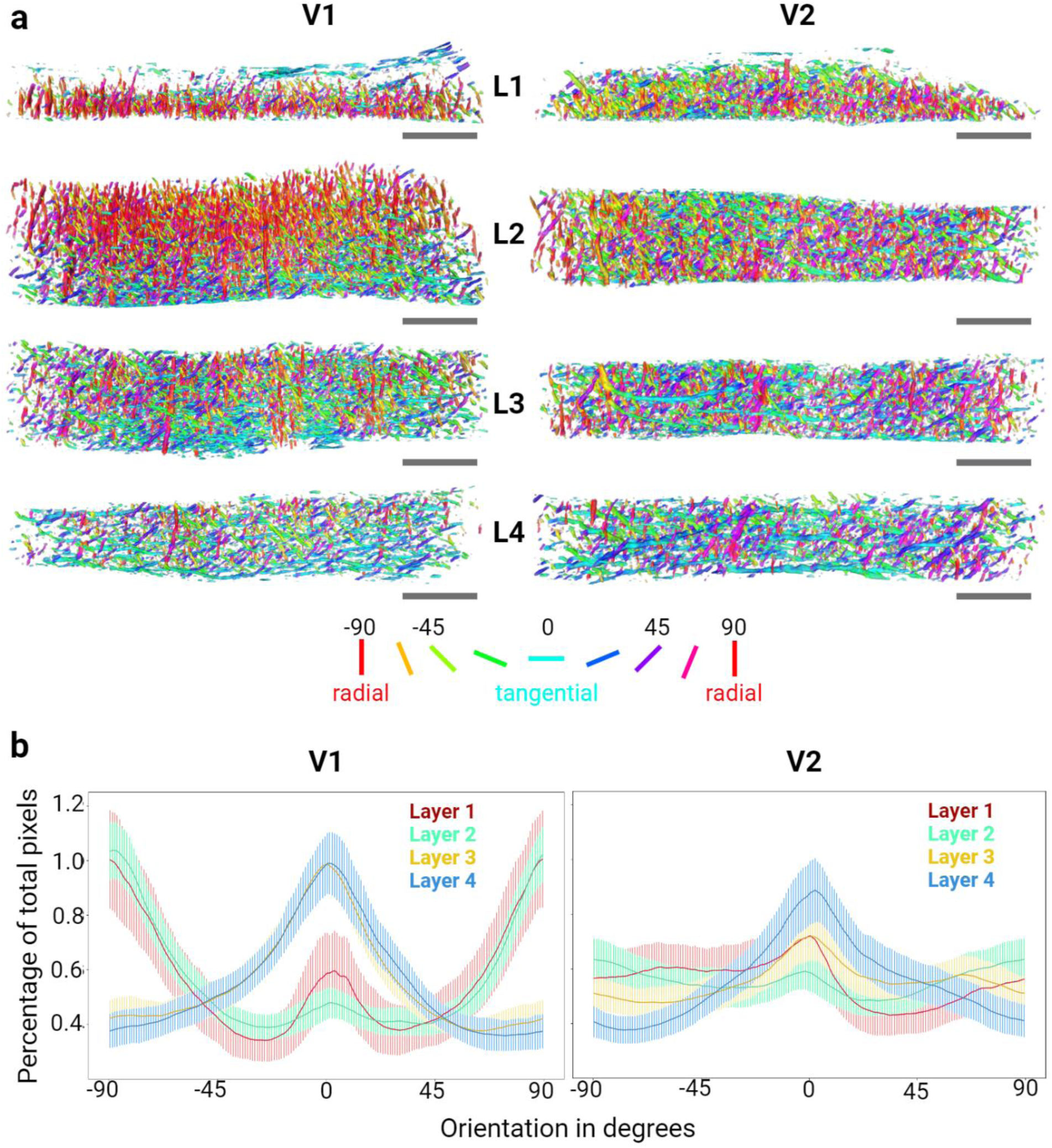
Vessel orientation profiles across angioarchitectural layers of human area V1 and V2. **a**) 3D renderings of the straightened angioarchitectonic layers (as delineated in supplementary figure 5). The renderings are colour-coded, with red corresponding to a 90° (vertical) orientation and cyan to a 0° (horizontal) orientation within the coronal plane. **b**) Vessel orientation profiles for each straightened layer over all images for V1 (left) and V2 (right). Shaded lines show the standard deviation. Scale bars: 400 µm respectively.

### 3.5 Vessel orientation profiles across infra-, supra- and granular compartments

One important aspect e.g., for layer fMRI studies, is the question of how angioarchitectural features correlate with the cytoarchitecture. Since even UHF MRI is not able to clearly delineate all 6 cytoarchitectural layers of the cortex, we decided to compare orientation profiles across the supragranular, granular, and infragranular compartments of the cortex. We will refer to these as functional compartments in the following (see *2.4.1 Orientation analysis*). As in the previous section, we show the results of the straightened layers in figure 8 to account for cortical curvature. The supragranular layer of V1 comprises both angioarchticture layers 1 and 2. Therefore, as one would expect, the main orientation in this layer is along the radial direction with a smaller second peak around the tangential direction. Interestingly, when dividing the volume into functional compartments, the granular layer shows a more even distribution of orientations, with a noticeable difference in the percentage of radially oriented vessels compared to the infragranular layer. The infragranular vessels are predominantly tangentially oriented, matching largely the distribution observed in angioarchitecture layer 4. As described for the angioarchitecture layers, the orientation profiles in the curved data volume are similar, but more equally distributed over all orientations (Suppl. Fig. 7). In the V2 dataset, the supragranular layer includes angioarchitecture layer 1, 2, and part of 3. In contrast to the orientation distributions between angioarchitectural layers, there is a clearly visible predominance of the radial directions in the supragranular layer, when compared to granular and infragranular layers. The profiles of granular and infragranular layers show substantial overlap, even though the granular layer is part of angio-layer 3, while the infragranular layer coincides almost completely with angio-layer 4. In both granular and infragranular layers, the majority of blood vessels oriented tangentially. However, with this parcellation scheme, there are discernible smaller peaks around the 45/-45° angles in the infragranular layer.

**Figure 8:**
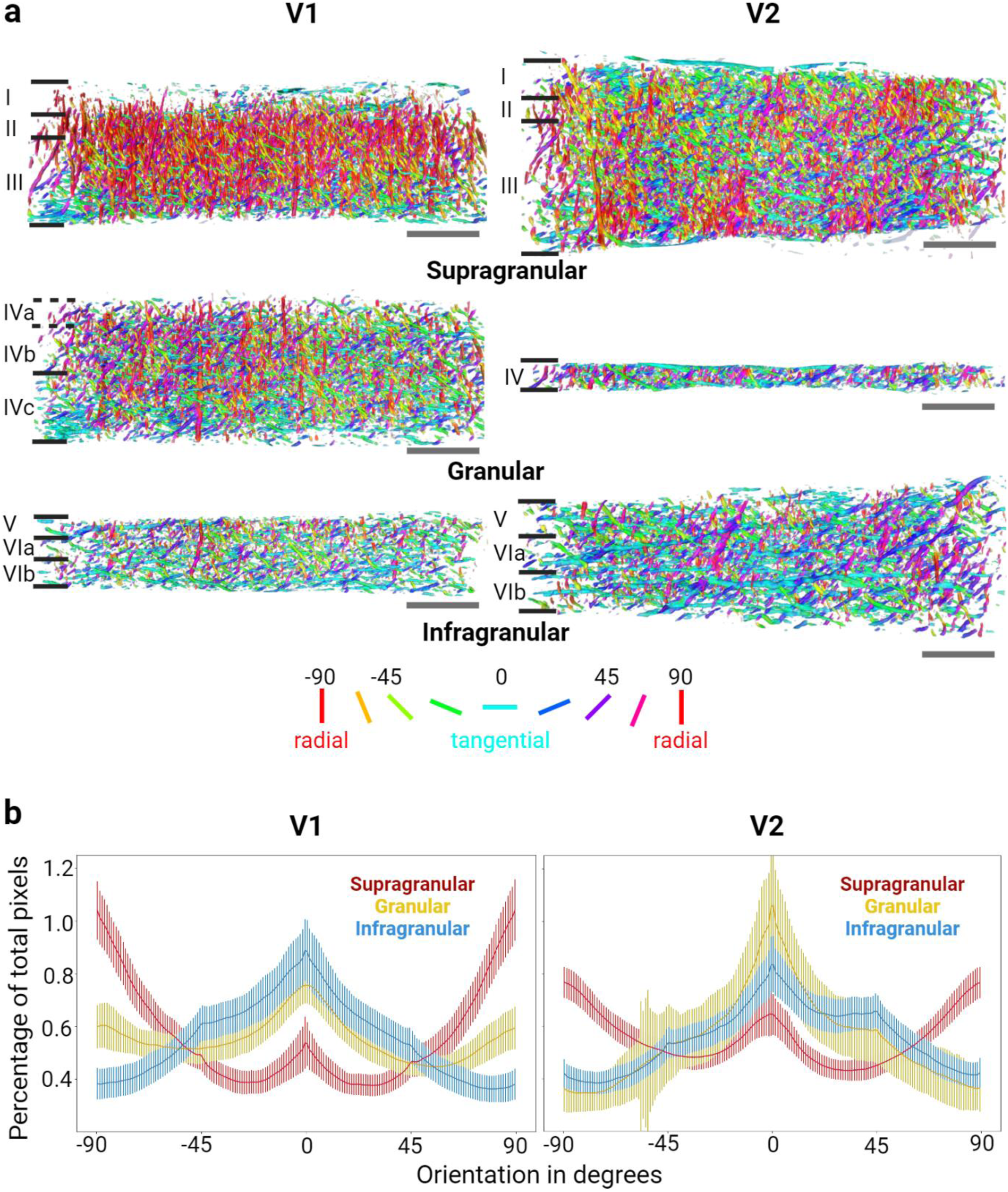
Vessel orientation profiles across infra-, supra- and granular compartments of human area V1 and V2. **a**) 3D renderings of the straightened functional compartments (as delineated in Supplementary figure 7). The renderings are colour-coded, with red corresponding to a 90° (vertical) orientation and cyan to a 0° (horizontal) orientation within the coronal plane. **b**) Vessel orientation profiles for each straightened layer over all images for V1 (left) and V2 (right). Shaded lines show the standard deviation. Scale bars (grey): 400 µm respectively.

Vessel orientation analysis over depth stratified by tangential and radial orientation reveals a general pattern across both areas for more radial blood vessel in supragranular and more tangential vessels in infragranular layers (Fig. 9). Close to 40 percent of the blood vessels in the most superficial layers of V1 have an orientation of 90° (±20°). In V2, this fraction makes up about a third of the blood vessels. Contrary to our expectations, the radially oriented vasculature, while more dominant, was not in the majority in either area in superficial layers. Overall, the distribution of radial vessels in both areas across cortical depth is similar and the relative contribution for both areas roughly the same, with exception of the most superficial part of the cortex, which is higher for V1. This is contrary to the qualitative visual impression of the data, but note that the graph shows relative differences for each area over depth, not absolute contributions of radial/tangential vasculature when comparing V1 and V2. Tangential vessels contribute approximately 10 percent to superficial layers and increase steadily towards over 60 percent close to the white matter. While this increase is sharper and more sudden in V1, the overall ratio of tangential vessels was found to be similar for both areas.

**Figure 9:**
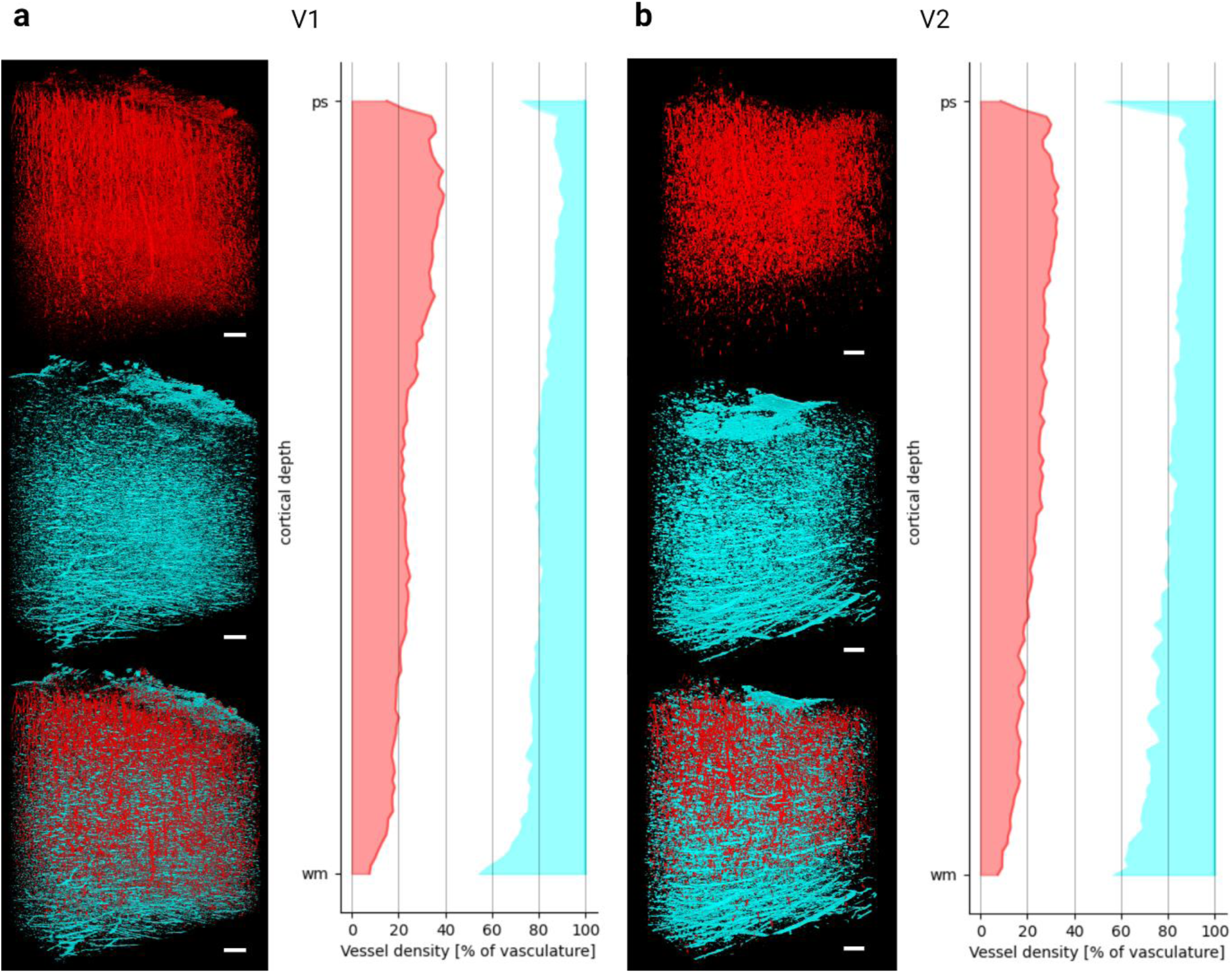
Vessel Orientation over depth stratified into radial and tangential vasculature. **a)** Radial and tangential vessels in V1. 3D renderings of the vasculature with (left) showing only radial (± 20°) vessels in red and tangential (± 20°) vessels in cyan. The graph shows the percentage of radial and tangential vasculature across cortical depth (right). **b)** Demonstrates the same for V2. Scale bars: 200 µm respectively.

A summary of the results for different angio- and cytoarchitectonic features, for each of the cytoarchitecture layers, is show in table 1 (V1) and table 2 (V2). For more details, see also the information provided in the supplementary tables. It is clear that while V1 has both an overall higher vascular and cell density as compared to V2, there is no clear correlation between the cell body and vessel density within each area, across layers.

**Table 1:** Summary statistics of cell and vascular densities for V1 across cytoarchitecture layers.

| layer | layer thickness [mm] | cell density [10<sup>3</sup> cells per mm<sup>3</sup>] | total vessel density [%] | radial vessel density [%] | tangential vessel density [%] |
| --- | --- | --- | --- | --- | --- |
| I | 0.211 | 41.6 | 6.6 | 2.2 | 0.9 |
| II | 0.102 | 114.4 | 8.7 | 3.2 | 1 |
| III | 0.466 | 106.1 | 10.1 | 2.8 | 1.7 |
| IVa | 0.09 | 81.6 | 11.5 | 2.5 | 2.5 |
| IVb | 0.166 | 64.6 | 12.8 | 3 | 2.5 |
| IVc | 0.252 | 76.2 | 12.6 | 2.8 | 2.6 |
| V | 0.113 | 65.3 | 10.4 | 2 | 2.4 |
| Vla | 0.088 | 77.2 | 10 | 1.8 | 2.4 |
| Vlb | 0.185 | 58 | 8.2 | 1.1 | 2.5 |

**Table 2:** Summary statistics of cell and vascular densities for V2 across cytoarchitecture layers.

| layer | layer thickness [mm] | cell density [10<sup>3</sup> cells per mm<sup>3</sup>] | total vessel density [%] | radial vessel density [%] | tangential vessel density [%] |
| --- | --- | --- | --- | --- | --- |
| I | 0.174 | 45.2 | 6.4 | 1.8 | 1 |
| II | 0.088 | 95.9 | 6.9 | 2.2 | 0.9 |
| III | 0.594 | 62.3 | 8.6 | 2.3 | 1.5 |
| IV | 0.164 | 73.1 | 8.6 | 1.8 | 1.9 |
| V | 0.183 | 46.3 | 8.6 | 1.5 | 2.2 |
| Vla | 0.128 | 42.5 | 7.8 | 1.3 | 2.2 |
| Vlb | 0.26 | 27.2 | 6.3 | 0.7 | 2.1 |

## Discussion

The angioMASH method described here enables the comprehensive study and three-dimensional mapping of angioarchitecture in formalin-fixed human cortical tissue. This work builds on and extends our earlier studies^41,71^ by combining cyto- and angioarchitecture labelling in the same sample, and by multi-color, multi-resolution imaging of the two labels in large FOVs. Moreover, we demonstrate quantitative data analysis for millimetre-thick volumetric datasets with the focus on intra- and inter-areal comparison of blood vessel orientation and densities. This combination of multiscale volumetric imaging of angio- and cytoarchitecture in the human cortex is, to the best of our knowledge, unique. For example, Harrison et al.^53^ and Zhao et al.^46^ processed and imaged human samples labelled for vasculature with light-sheet and confocal microscopy, but show no analysis of the angioarchitecture. Here, we focussed on samples of an intermediate size (∼30 mm x 10 mm x 3 mm), as the imaging of much thicker samples quickly becomes more challenging, mostly due to light scattering, even with light-sheet microscopy and optical tissue clearing.

Our findings match well with the descriptions of human and non-human primate visual cortex by other authors, which were based on the imaging of thin 2D sections^5,31,32^, lending support to the high labelling quality achievable with the angioMASH technique, despite the *en bloc* labelling of millimetre thick samples. The density distribution found in V1 of the current samples is highly similar to that in other reported primate species^4,32^ with a noticeable high density in layers IVb-c. In V2 our observed distribution matches the general description of Duvernoy^5^, who remarked that lower (cytoarchitectural) layer III shows at least as high vascular density as layer IV. In our data, the vascular density of layers III and IV were identical, matching his description. This contrasts with Weber’s observation in the macaque cortex, in which the highest density being located in layer IV^32^. Future multispecies investigations could determine if this finding represents an inter-species difference. Likewise, it remains to be determined whether other areas than V1 and V2 in the human brain are different in their layered angioarchitecture or if the difference is mainly found in the comparison of primary sensory areas (such as V1) with other areas (beginning with V2)^31^.

Human cortical angioarchitecture is currently not extensively researched with 3D microscopy, while it has become of increasing importance for modern non-invasive imaging techniques whose signal is strongly coupled to the underlying vasculature and hemodynamics, such as fNIRS^18,19^ and fMRI^20,21^. The data and results presented here provide a first ground truth for some relevant vascular parameters for these non-invasive imaging methods at a meso- and microscopic level. Here, the variability of relative vascular density over cortical depth, vessel orientation within cortex, and intra-areal variation over V1 and V2, are of particular interest. Based on the results presented here, as well as data by other groups, a direct one to one relationship between cyto- and angioarchitecture does not seem to exist^30–32,72^. However, it has been shown that the overall densities of cells and vasculature are correlated when comparing interareal averages^72^. Our density comparison shows that there is higher vascular density in the supragranular and granular layers in V1 as compared to V2, while the infragranular vessel density is similar for both areas. The density curves across cortical depth for V1 and V2, with increasing vasculature towards the middle layers, and decreasing density towards infragranular layers, seem to be a more general feature of angioarchitecture across different areas of the human brain^30^ as well as across species^31^. From this it seems to follow, that primary sensory cortices demonstrate their highest vasculature around their extensive input layers, whereas other (higher) areas show at least equally high vascular density in supragranular compared to granular layers. This might reflect metabolically expensive internal computations represented at various cortical depth, dependent on the precise function of the brain area^73^. It would be intriguing to compare more areas including higher-level association areas as well as primary motor cortex, to see if this simple model of vascular density following the metabolic demands of the respective cortical module holds true.

### Technical considerations

A distinction between the different components of the vascular network into arteries, arterioles, veins, venules, and capillaries is not possible with the current angioMASH protocol. The tomato lectin applied here as a vessel label, binds to specific sugar-groups present in the endothelial cells of the vessel wall. Therefore, all the blood vessels, including the capillaries are labelled. However, distinctions based on morphology might be possible. For example, based on relative vessel diameters, arteries and veins might be distinguished from arterioles and venules as well as capillaries. Alternatively, other labelling approaches might also be used. Recently, the Renier laboratory published an outstanding atlas of the mouse brain vasculature, based on an elaborate staining procedure with several clearing compatible antibody labels to distinguish arterial and veinous components^50^. A different approach was taken by the Ertürk group, which used dye injections and machine learning to differentiate between the components in the cleared mouse brain^52^ and similar approaches have been used by other groups as well^74,75^. For future studies, the adaptation of one of the two approaches might allow a more detailed description of the vasculature in the human brain. If antibodies against vein- or artery-specific epitopes could be introduced deep enough into the much denser human brain tissue, these components could be separated into different channels.

A disadvantage of both lectin and antibody labels is that they only stain the vessel wall. This is problematic because it can lead to faulty segmentation of large vessels into two structures^50^, depending on the angle at which the imaging plane intersects the vessel. In addition, larger blood vessels often collapse during the histological processing, which makes vessel diameter estimates challenging. We tried to account for this by using the perivascular spaces visible as a negative image in the cytoarchitecture channel for diameter estimates of the larger vessels. However, this might slightly overestimate these diameters. Injection techniques have the advantage of filling the entire lumen, which should prevent larger vessels from collapsing during the clearing. This method would have the additional advantage of increasing the signal-to-noise ratio, making deep imaging easier^74^. A traditional approach to label angioarchitecture in the human brain, is the injection with Indian ink gel^5,30,40^. Adapting the classic Indian ink protocol towards a fluorescent, clearing compatible label, could allow for much better imaging deep into human brain samples and would allow for a more reliable determination of the diameters of larger vessels. One potential limitation of these injection-based techniques, specifically in the human brain, are blood coagulations in the smallest vessels due to *post mortem* delays, which might lead to them being insufficiently labelled. Here, the combination with angioMASH could prove particularly useful.

Another limitation of the current study is that the orientation analysis, was carried out on 2D planes across the 3D volume rather than directly in 3D. In the future, a more sophisticated 3D analysis of the datasets acquired in this study could reveal more extensive spatial relationships. For instance, some orientations might be better represented in a 3D vascular skeleton, instead of the current 2D projection. The current orientation analysis makes use of an existing function in FIJI to straighten and remove the slight curvature in the 2D projections. In our coronal sections, it is then assumed that the vertical orientation corresponds to the radial direction in the cortex and the horizontal axis to the tangential direction. Unfortunately, this aspect of the pipeline does not scale well to larger datasets, which include entire gyri and have more pronounced curvature of the cortical ribbon. Given the complex curvature of the human cortex, it is also not always possible to guarantee a perfectly coronal sectioning plane. Establishing a cortical segmentation routine, which translates the dataset into a cortex-based coordinate system with radial, tangential, and oblique directions would be necessary to scale the analysis up to larger FOVs.

### Outlook

It has been shown here that the combined investigation of angio- and cytoarchitecture is possible and it is expected that the labelling and imaging routines will be scaled up in the near future to allow for the processing and analysis of much larger samples with angioMASH^71^. We have recently shown this kind of upscaling for our cytoMASH pipeline^71^ and deem it feasible, therefore, to extend angioMASH in the same way. This kind of upscaling is only considered possible with an oblique light-sheet set-up^71,76,77^, which has the advantage of enabling a more homogenous imaging of very large, flat samples such as whole brain slices^71^.

The current datasets were acquired from the occipital lobes of healthy body donors. Going forward, two obvious next goals would be 1) the extension towards more brain areas, to get a more comprehensive overview of the healthy human angioarchitecture and 2) the comparison with pathological samples. Multiple brain disorders, such as vascular dementia^78^ or epilepsy^79,80^ are known to result in abnormal vascular organisation or changes in the vessel anatomy itself. angioMASH could enable the study of these disorders in the human brain in its full three-dimensional aspects. Optical tissue clearing and light-sheet microscopy are already starting to be used for 3D investigation of healthy and pathological human brain anatomy^43,44,46,81,82^. Taking histological investigations into 3D can help highlight features, which would be difficult to detect on conventional 2D sections, as has been shown for the spatial distributions of plaques in Alzheimer’s disease^44^.

Although large scale brain mapping endeavours using 3D reconstructions of 2D sections have yielded astounding results^83,84^, the combination of tissue clearing and light-sheet microscopy has the potential enable smaller laboratories to engage in similar brain mapping endeavours, with less time-investment and manpower. We predict that the further developments in 3D histology techniques will enable the study of brain anatomy in an unparalleled manner.

## Supporting information

Supplementary_tables_angioMASH

Supplementary_figures_angioMASH

Supplementary_video _1_angioMASH

Supplementary_video _2_angioMASH

Supplementary_video _3_angioMASH

Supplementary_video _4_angioMASH

## Author contribution

AR, MC, and SH came up with the experimental design. The angioMASH pipeline was designed and samples were created by SH. The tissue for this project was provided by AH. Imaging was performed by PB and SH on the infrastructure provided by FH and PB. Data analysis was primarily performed by JF, HH, and MC. Data vizualizations were generated by JF, HH, and SH. All authors have contributed in writing the manuscript.

## Acknowledgements

We would like to thank Kathleen Rockland for her invaluable input on the manuscript and her extremely generous time allocation for the delineation of the cytoarchitecture layers in our dataset. We would further like to thank our diligent student assistants for providing a helping hand in the lab. Without the help of many talented and laborious students, such as Mathilde Bertrand, George Burchell, and Jana Totzek, the method development would not have been possible.

## Notes

### Competing Interest Statement

The authors have declared no competing interest.

### Summary of Updates

New tables have been added in the manuscript and supplementary material. Figures 5 and 9 have been updated. Several minor errors in the manuscript text have been mended.

