## Supplementary_figures_angioMASH for "Investigating microscopic angioarchitecture in the human visual cortex in 3D with angioMASH tissue clearing and labelling"

### Supplementary material

**Supplementary Table 1: Patient information of the occipital lobe specimen.**

| Sample | Gender | Age | Prior neuropathological disease |
|---|---|---|---|
| Occipital lobe 1 | male | 82 | no |
| Occipital lobe 2 | female | 101 | no |
| Occipital lobe 3 | male | 90 | no |

**Supplementary Video 1.** 3D volume rendering of an overview acquisition of occipital lobe 3, sample 2. The rendering shows the penetration of the vessel label throughout the sample (resolution ~6 µm).

**Supplementary Video 2.** 3D volume rendering of the stitched high-resolution (~ 2 µm) raw data of occipital lobe 3, sample 2, showing the extent of the acquired tiled ROI.

**Supplementary Video 3.** 3D volume rendering of the orientation-coded blood vessels in V1 (red corresponding to radial and cyan to tangential vessels).

**Supplementary Video 4.** 3D volume rendering of the orientation-coded blood vessels in V2 (red corresponding to radial and cyan to tangential vessels).

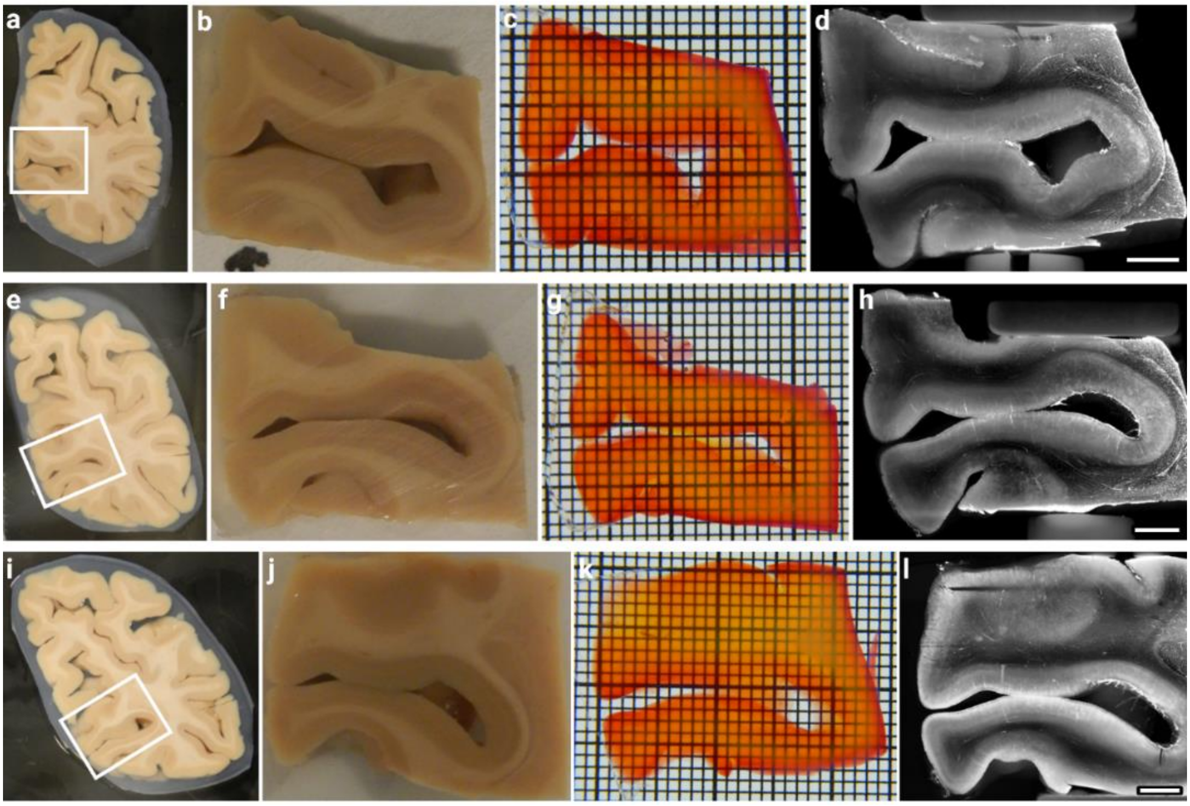


**Supplementary Figure 1: Overview of Occipital lobe 1 samples.** Three consecutive coronal slices (**a**, **e**, and **I**; direction posterior to anterior) where cut at a thickness of approx. 3 mm from the occipital lobe and the two gyri around the calcarine sulcus were blocked (**b**, **f**, and **j**). Blocked samples contained the V1/V2 border and parts of V2 in all cases. Samples show high transparency after labelling and clearing, especially in the ROIs of the gyri crowns (**c**, **g**, and **k**; grid size: 1x1 mm). Overview acquisitions with the highest possible FOV and a resolution of approx. 10 µm isotropic of the vessel label (**d**, **h**, and **l**; scale bars: 3 mm respectively). Already at this resolution, the V1/V2 border is visible (see especially panel **l**).

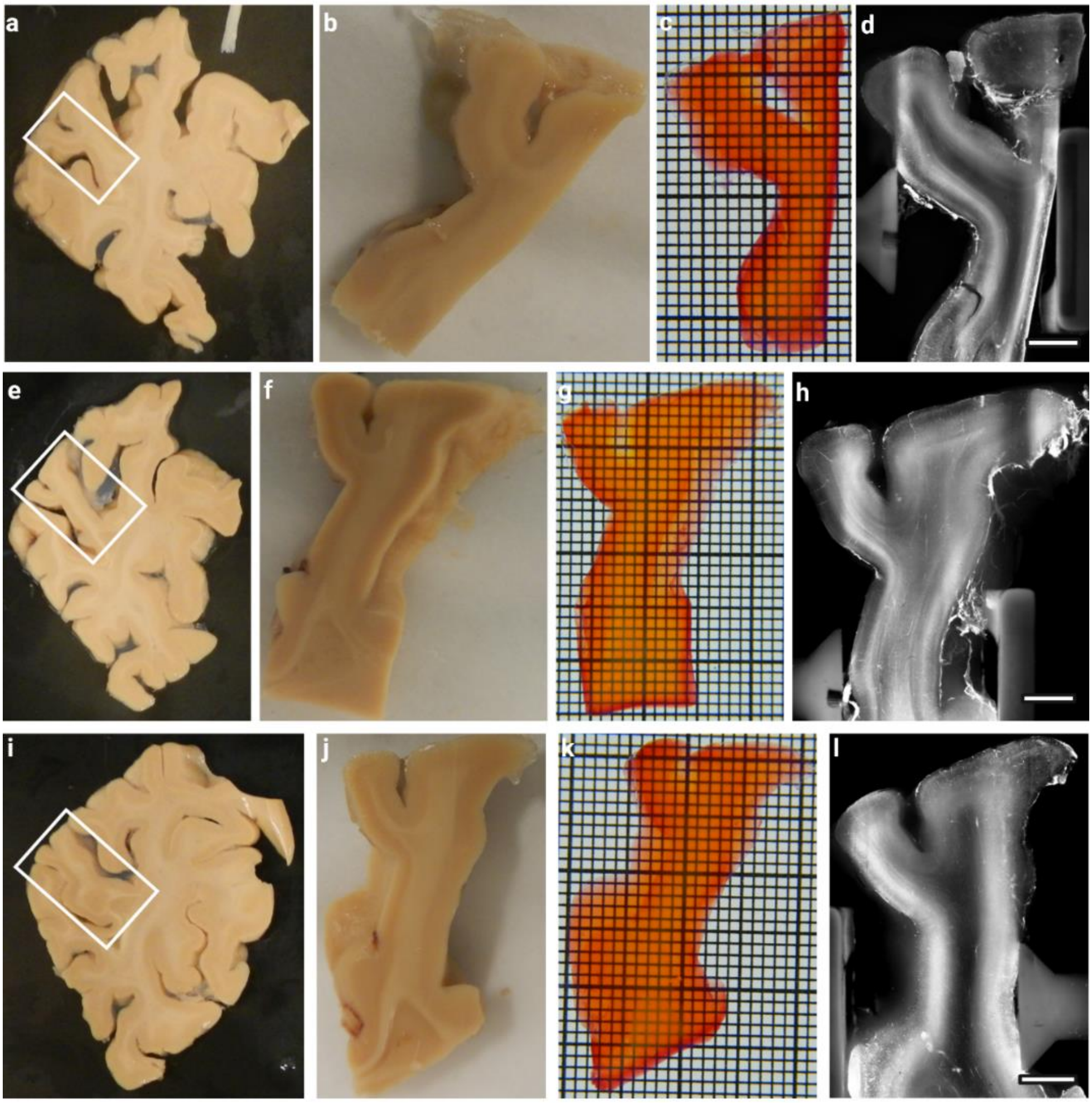


**Supplementary Figure 2: Overview of Occipital lobe 2 samples.** Three consecutive coronal slices (**a**, **e**, and **I**; direction posterior to anterior) where cut at a thickness of approx. 3 mm from the occipital lobe and samples containing a gyrus next to the calcarine sulcus were blocked (**b**, **f**, and **j**). Blocked samples contained the V1/V2 border and parts of V2 in all cases. Samples show high transparency after labelling and clearing, especially in the ROIs of the gyri crowns (**c**, **g**, and **k**; grid size: 1x1 mm). Overview acquisitions with the highest possible FOV and a resolution of approx. 10 µm isotropic of the vessel label (**d**, **h**, and **l**; scale bars: 3 mm respectively). Already at this resolution, the V1/V2 border is visible (see especially panel **h**).

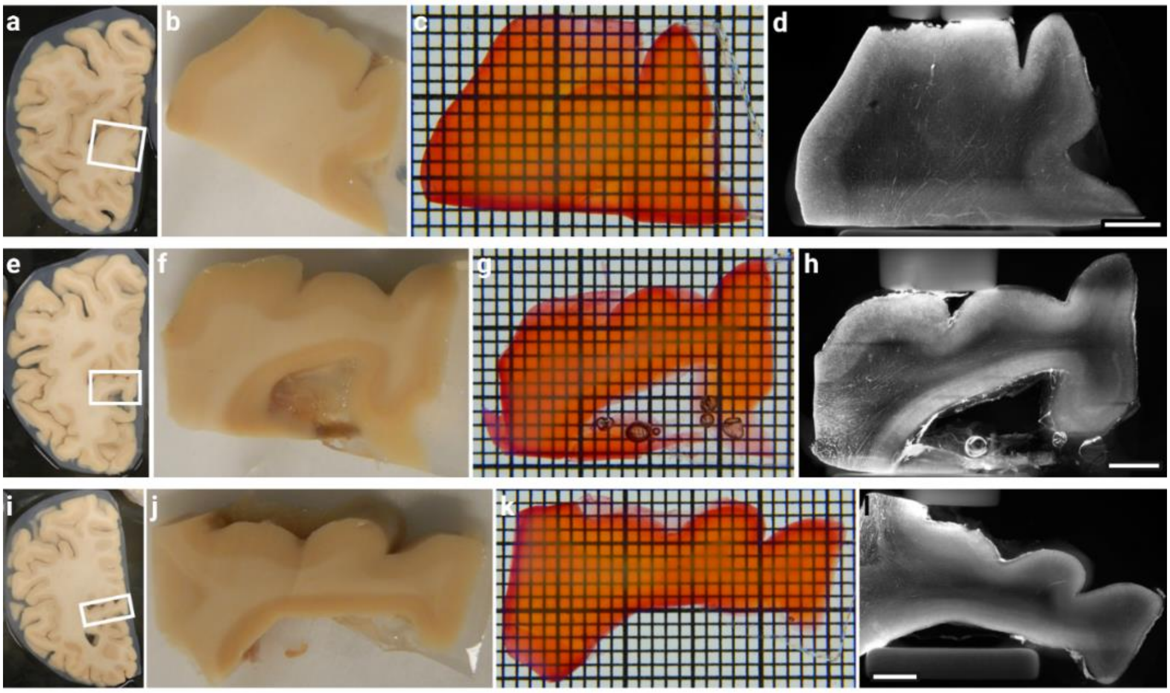

**Supplementary Figure 3: Overview of Occipital lobe 3 samples.** Three consecutive coronal slices (**a**, **e**, and **I**; direction posterior to anterior) where cut at a thickness of approx. 3 mm from the occipital lobe and samples containing a gyrus next to the calcarine sulcus were blocked (**b**, **f**, and **j**). Blocked samples contained the V1/V2 border and parts of V2 in all cases. Samples show high transparency after labelling and clearing, especially in the ROIs of the gyri corwns (**c**, **g**, and **k**; grid size: 1x1 mm). Overview acquisitions with the highest possible FOV and a resolution of approx. 10 µm isotropic of the vessel label (**d**, **h**, and **l**; scale bars: 3 mm respectively). Already at this resolution, the V1/V2 border is visible (see especially panel **h**).

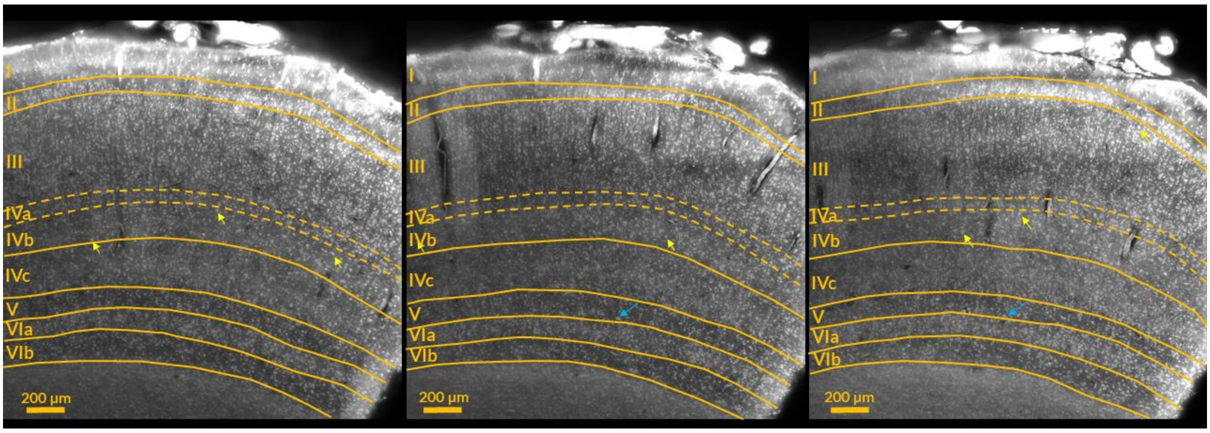


**Supplementary figure 4: Delineation of cytoarchitecture layers in V1.** Shown are 3 MIP over 30 µm of the same dataset, taken at different depth. Two expert annotators independently drew the layers in each MIP and reached a consensus thereafter. Layering was highly consistent between the different MIPs as well as between annotators, with the exception of layer IVa, which borders are therefore indicated tentatively by dashed lines. Yellow arrow indicate large cells in IVb, which were taken as an identifying criterion and blue arrows show Meynert cells at the border of layers V and VI.

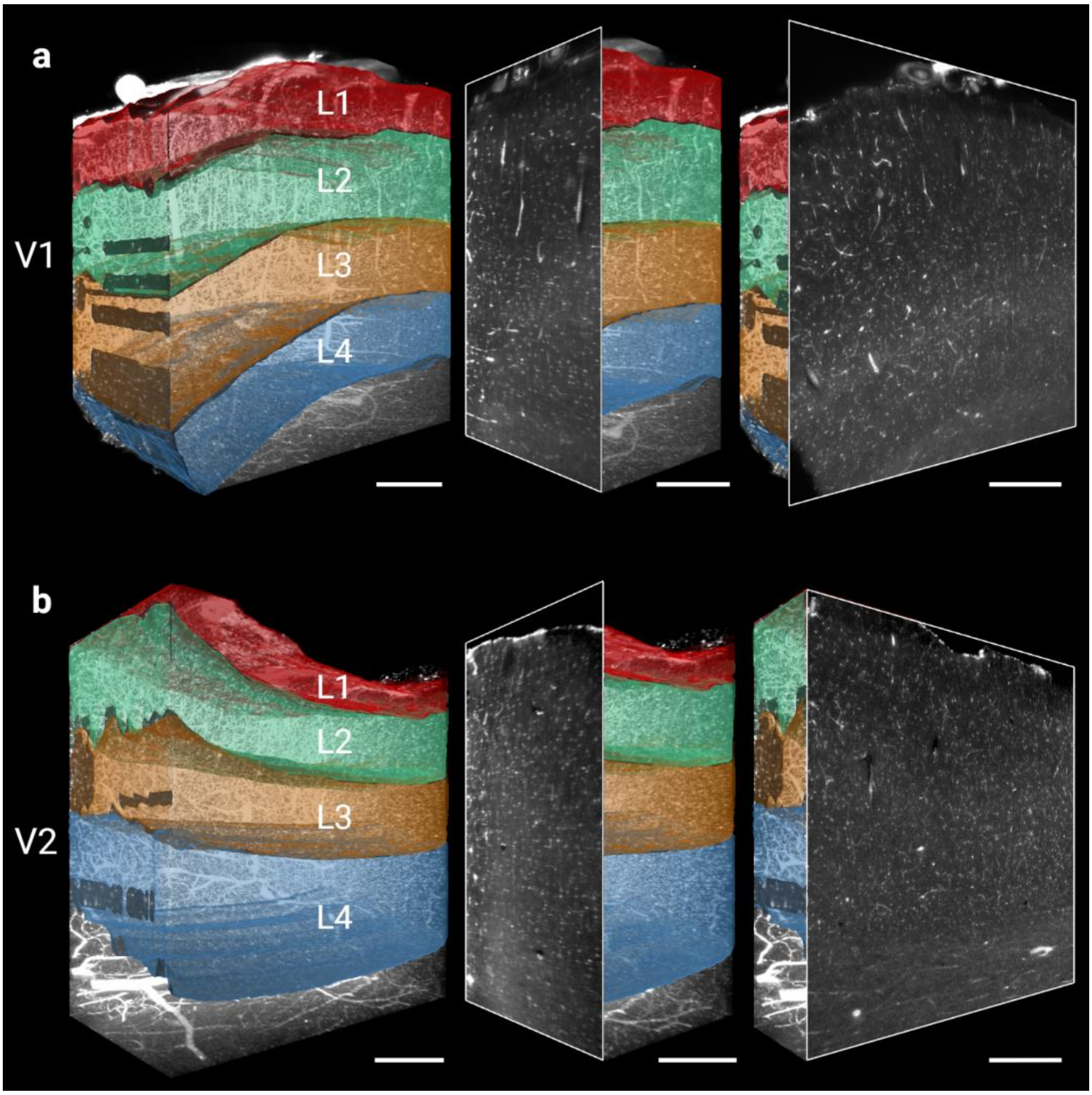


**Supplementary figure 5: Visualisation of the four main angioarchitectonic layers according to Duvernoy. a**) Delineation of the four main vascular layers in the V1 dataset. A rendering of the whole volume (left) is complemented by a view onto the YZ-plane (middle) and the XY-plane (right). **b**) Showing the same as in (**a**) for V2. Note the different distribution of the corresponding layers (e.g. layer 1 and 4 in V1 compared to V2). The darker areas on the left side on some of the layers are artefacts, which occurred when the manual layer segmentation was a few pixels away from the edge of the image. Scale bars: 400 µm respectively.

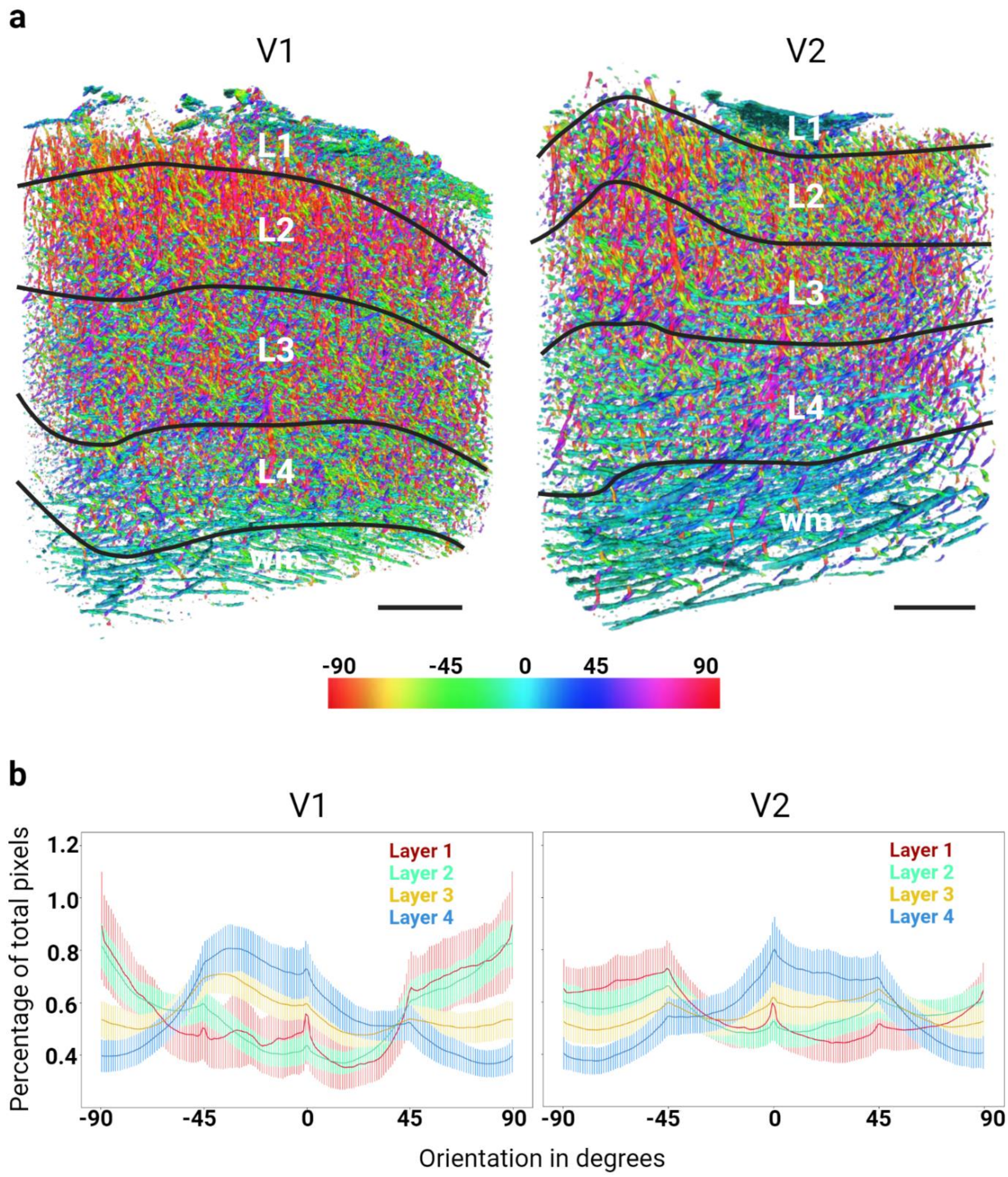


**Supplementary figure 6: 3D rendering of planar vessel orientation in human area V1 and V2. a**) 3D rendering of the colour-coded blood vessel for V1 (left) and V2 (right). Red corresponds to vertically oriented vessels and cyan to horizontal blood vessels in the coronal plane (XY-plane of the sample). The volume had been divided into the four angioarchitecture layers according to Duvernoy's definition (black lines; wm = white matter). **b**) Mean distributions of the relative amount of pixel orientations in degrees for each layer over all images for V1 (left) and V2 (right). Shading depicts the standard deviation. Scale bars: 400 µm respectively.

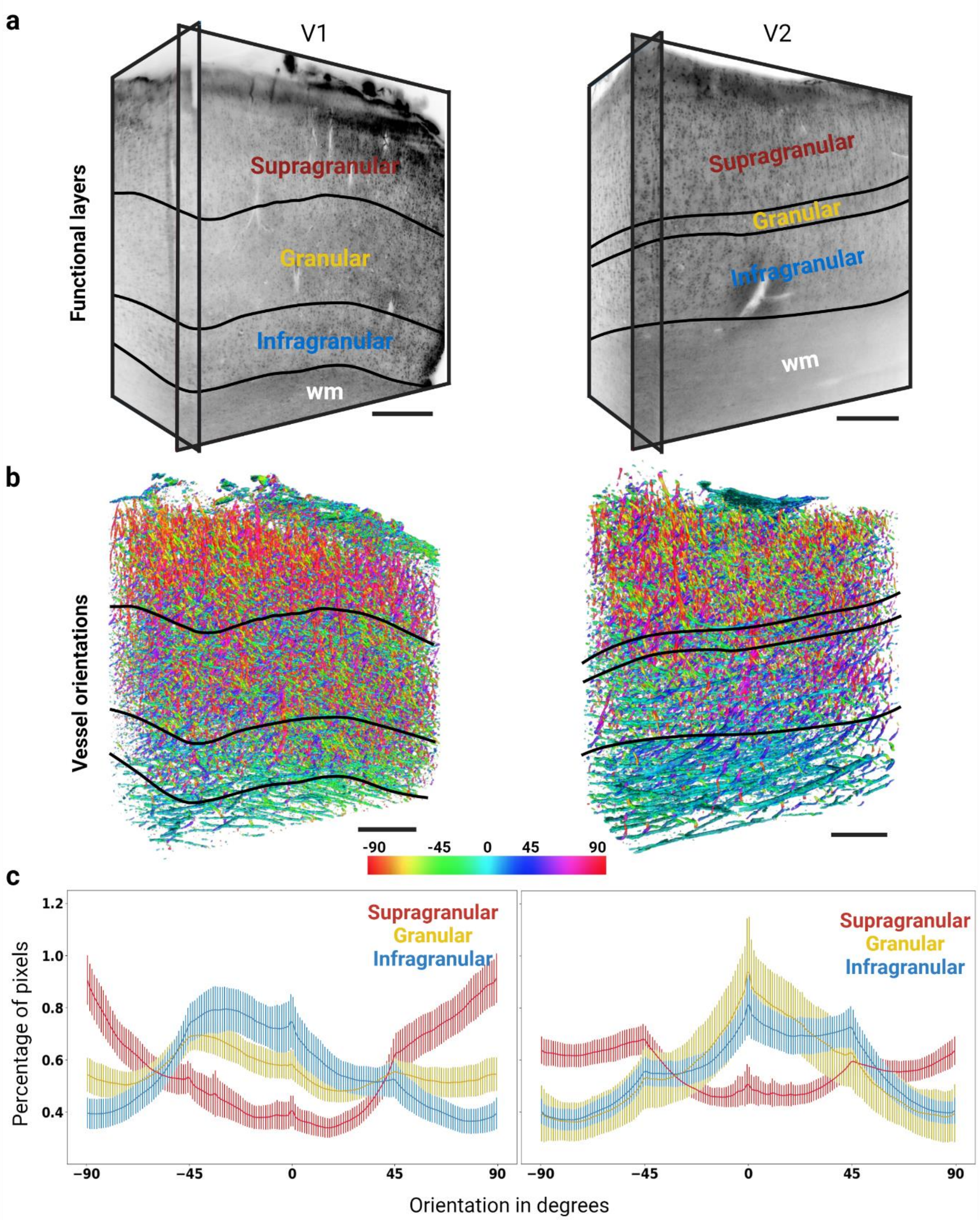


**Supplementary figure 7: Distribution of planar vessel orientation in human area V1 and V2, in relation to the cytoarchitecture. a**) 3D volume of the cytoarchitecture with the MASH-NR cell body label for V1 (left) and V2 (right). For better visualisation of the extremely dense label, two orthogonal planes are shown, rather than a rendering (black frames). The lines indicate the supragranular, granular, and infragranular cytoarchitectonic compartments, respectively. **b**) 3D renderings of blood vessel orientation for V1 (left) and V2 (right). Black lines indicate the same compartments that have been explained in **a**. **c**) Mean distributions of the relative amount of pixel orientations in degrees for each layer over all images for V1 (left) and V2 (right). Shading depicts the standard deviation. Scale bars: 400 µm respectively. wm = white matter.
